# Distinguishing new from persistent infections at the strain level using longitudinal genotyping data

**DOI:** 10.1101/2025.02.06.636982

**Authors:** William A. Nickols, Philipp Schwabl, Amadou Niangaly, Sean C. Murphy, Peter D. Crompton, Daniel E. Neafsey

## Abstract

**Motivation:** Longitudinal pathogen genotyping data from individual hosts can uncover strain-specific infection dynamics and their relationships to disease and intervention, especially in the malaria field. An important use case involves distinguishing newly incident from pre-existing (persistent) strains, but implementation faces statistical challenges relating to individual samples containing multiple strains, strains sharing alleles, and markers dropping out stochastically during the genotyping process. Current approaches to distinguish new versus persistent strains therefore rely primarily on simple rules that consider only the time since alleles were last observed.

**Results:** We developed DINEMITES (**Di**stinguishing **Ne**w **M**alaria **I**nfections in **T**im**e S**eries), a set of statistical methods to estimate, from longitudinal genotyping data, the probability each sequenced allele represents a new infection harboring that allele, the total molecular force of infection (molFOI, the cumulative number of newly acquired strains over time) for each individual, and the total number of new infection events for each individual. DINEMITES can handle time points with missing sequencing data, incorporate treatment history and covariates affecting the rate of new or persistent infections, and can scale to studies with thousands of samples sequenced across multiple loci containing hundreds of possible alleles. In synthetic evaluations, the DINEMITES Bayesian model, which generally outperformed an alternative clustering-based model also developed in this work, accurately estimated key clinical parameters such as molFOI (bias 2.5, compared to −12.2 for a typical simple rule). When applied to three real longitudinal genotyping datasets, the model detected 33%, 112%, and 359% more average infections per participant than would have been detected by applying a typical simple rule to the equivalent datasets without sequencing.

**Availability and implementation:** DINEMITES is freely available as an R package, along with documentation, tutorials, and example data, at https://github.com/WillNickols/dinemites.

## Introduction

Pathogen genetic analysis is key to understanding infectious disease dynamics, especially for diseases such as malaria in which the etiological agent (*Plasmodium* spp.) comprises multiple strains which can co-infect individual hosts. In the malaria field, cross-sectional population genetic analyses are frequently used to track *Plasmodium* parasite population connectivity, identify polymorphisms linked to therapeutic resistance, and monitor the dispersal of such variants.^1,2,3,4^ There is also interest in the potential for longitudinal genotypic analyses to elucidate within-host malaria infection dynamics, especially in the context of intervention efficacy trials.^5,6,7,8,9^ Such trials sample participants serially over the course of months to years to evaluate whether the intervention (e.g., a monoclonal antibody or vaccine) effectively blocks new infections.

Genotypic data have the potential to be particularly informative for longitudinal infectious disease studies because they can both inform how many unique pathogen strains were present during an infection and distinguish new from pre-existing or recrudescent infections, neither of which is possible with binary (presence/absence) infection detection methods such as microscopy or (q)PCR. However, several methodological and statistical challenges remain to be addressed to obtain the maximum information available from such data for pathogens such as *Plasmodium*.

One challenge for interpreting longitudinal *Plasmodium* genotyping data is presented by polyclonal infections, in which two or more genetically distinct strains simultaneously infect a host. The complexity of infection (COI; the number of distinct co-infecting strains) is often epidemiologically important but estimating COI is complicated by factors such as partial genetic relatedness of co-infecting parasite lineages and the choice of which loci to sequence, among other factors.^10,11,12^ Previous methods have been developed to deconvolve infections from independent samples (e.g., studies with one sample per individual) to estimate COI,^13,14,15^ but longitudinal studies clearly involve non-independent samples because the same individuals are sampled repeatedly over time.

Whether infections are monoclonal or polyclonal, a statistically informed approach is needed to distinguish new from persistent strain infections. This capacity would inform malaria intervention studies, where the efficacy could be measured as cumulative protection against infection over time rather than simply time to the first infection or case, or via other metrics such as strain-level asymptomatic infection duration.^5,16,17,18^ However, this task presents statistical challenges since alleles observed repeatedly can come from either persistent infections or new infections with the same allele. Furthermore, alleles can stochastically be missed during sequencing due to low pathogen abundance, even when an infection is detectable using more sensitive molecular methods such as ultra-sensitive qRT-PCR.^19^ Historically, studies have typically identified new infections from such data using simple evidence-informed rules based on the time since the last detected infection.^5,6,8,20,21^ The few existing statistical methods addressing this application are limited by the fact that they address new infections only at an aggregate level rather than the strain level; only analyze a small number of time points per subject; do not account for the challenge of detection-positive but sequencing-absent samples; require entire genomes; or are locus- or study-specific.^22,23,24^

To bridge this gap, we introduce DINEMITES (**Di**stinguishing **Ne**w **M**alaria **I**nfections in **T**im**e S**eries) to estimate, from longitudinal genotyping data, (1) the probability each sequenced allele represents a new infection with that allele, (2) the molecular force of infection for each individual (molFOI, the cumulative number of newly acquired strains over time), and (3) the total number of new infection events for each individual. Additionally, DINEMITES handles detection-positive (e.g., via qPCR) but sequencing-absent time points and scales to studies involving hundreds of participants with dozens of time points each, sequenced across multiple loci containing hundreds of possible alleles. DINEMITES, along with its source code, documentation, tutorials, and example data sets, is freely available at https://github.com/WillNickols/dinemites.

## Results

Here, we present DINEMITES, a set of computational tools for analyzing longitudinal infection genotyping data. DINEMITES takes as input the pathogen genotyping results for each individual at each time point, along with (optionally) a record of samples which are known to be pathogen-positive but lack sequencing data, a record of when individuals were treated for the pathogen, and other infection-modifying covariates such as season, insecticide net usage, and vaccination status (**Fig. 1A**). The software outputs the probability each sequenced allele belongs to a new infection, the per-individual molFOI, the per-individual number of new infection events, and plots visualizing the time series for each individual.

**Figure 1.**
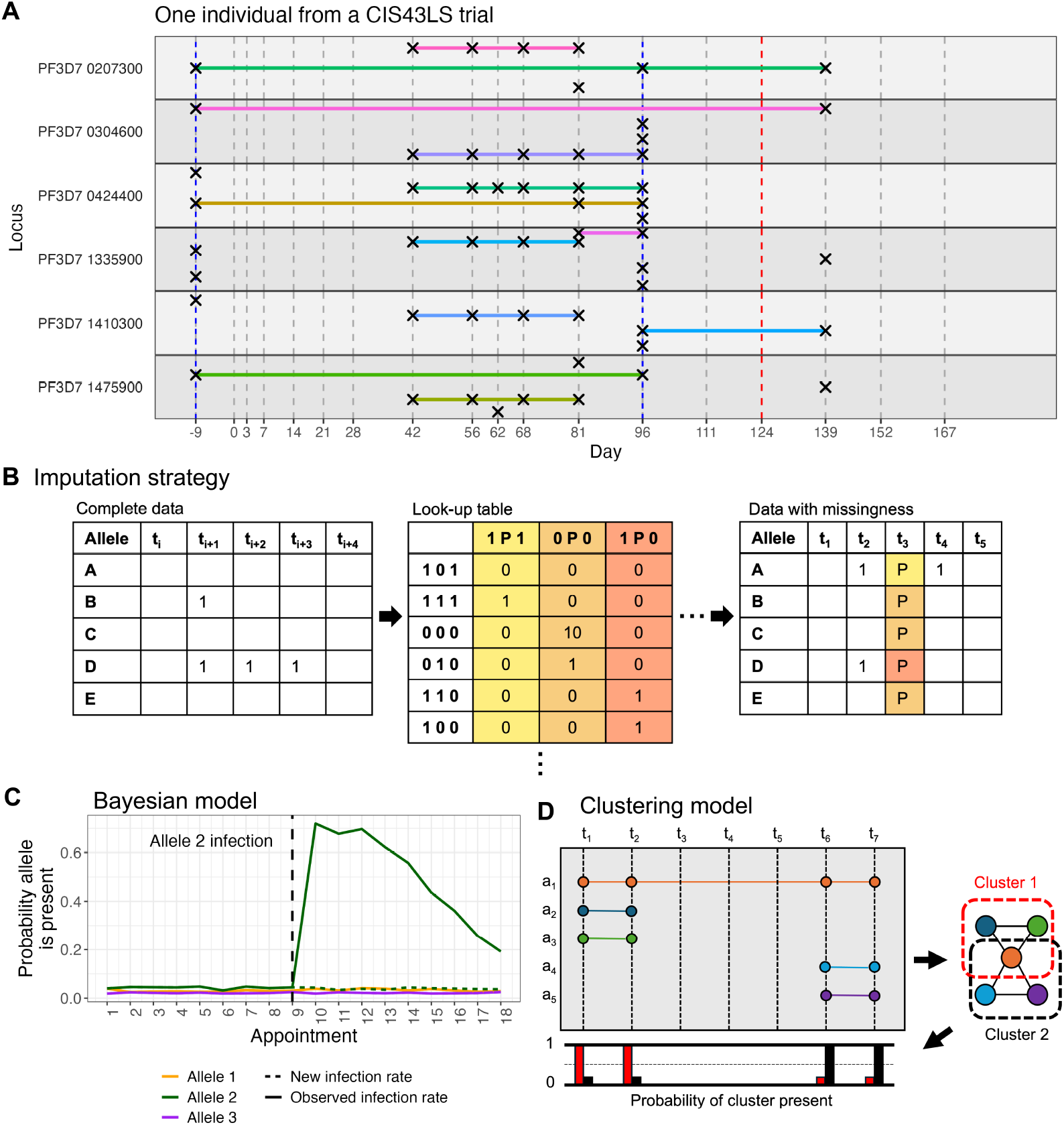
Statistical methods distinguish alleles from new versus persistent infections in longitudinal genotyping data. **A**. Longitudinal genotyping data consist of multiple time points (vertical dashed lines) at which subjects provided blood spots. From these, multiple loci (horizontal panes) are sequenced to detect alleles (crosses if present), sometimes occurring repeatedly over time (indicated by horizontal lines). Some time points (e.g., day 96) involve polyclonal infections, but not all loci have the same number of alleles. Some time points are only known to be pathogen-positive by qRT-PCR but have no genotyping information (red dashes, day 124). Participants are sometimes treated for the pathogen (blue dashes). This example is from subject 0266R of a CIS43LS monoclonal antibody study.^16^ **B**. Some time points are only known to be positive (P) for an infection but do not have genotyping information (missingness). The multiple imputation procedure uses the available data with genotyping (alleles displayed as 1 if present, blank if absent) to build a table of how frequently each missingness pattern could be generated by the complete data and uses this to impute the missing data. **C**. The Bayesian model determines the probability an allele is new by comparing the probability of observing the allele due to a new infection to the probability of observing the allele at all, the latter being increased after a previous detection. **D**. The clustering model creates a graph from co-occurrence of the alleles, clusters the edges to infer infection events, performs likelihood maximization to estimate the probability each cluster was present on each day, and computes the probability an allele was new based on how likely it was to be present in each cluster and how likely each cluster was to be new on each day.

In some studies, a subset of samples are known to be pathogen-positive but do not have genotyping data, either because a more sensitive detection method was used alongside genotyping (e.g., qPCR or qRT-PCR^5,6,16,19^) or because only some samples were chosen for genotyping (e.g., to reduce costs). Such samples are hereafter referred to as missing (distinct from samples for which no data are available). To handle such samples, a multiple imputation procedure^25^ uses the portion of the data with complete genotyping information to build a table of how often each missingness pattern could be generated by the complete data (**Fig. 1B, Methods**). These frequencies are then used to sample specific alleles at the missing time points, creating multiple complete datasets which are analyzed uniformly but separately, from which the results are ultimately aggregated. To determine whether each sequenced allele is new in a complete or imputed dataset, one of three methods is applied: (1) a simple rule that defines an allele as persistent if it has been recently observed, (2) a Bayesian model that compares the probability of a new infection by that allele to the probability of observing that allele after a prior infection (**Fig. 1C**), or (3) a clustering model that uses allele co-occurrence to define likely infection events (**Fig. 1D, Methods**).

### A Bayesian model with multiple imputation outperforms other models on synthetic data

To evaluate the ability of DINEMITES to accurately classify infections as new or persistent, synthetic longitudinal genotyping datasets were generated in which each allele was known to come from a new or persistent infection. The genotyping data were generated according to either a model with probabilities of infections that updated at each time point based on the previous time points (“rolling presence probability”) or a model with infections lasting for variable time periods sampled from a Poisson distribution (“Poisson time to clearance”) (**Supplementary Information**). These datasets were created to simulate high-transmission settings with common occurrences of superinfection (new infection in the presence of persistent infection) and polyclonality (multiple alleles present at a single locus, which corresponds to multiple strains occurring because *Plasmodium* parasites are haploid in the blood stage). To mimic missingness in real genotyping data, varying proportions of the genotyping-positive time points were masked so that they were only known to be infection-positive (qPCR positive). Then, with and without the imputation strategy, the Bayesian model, the clustering model, and a simple rule were applied to the datasets to determine the probability each sequenced allele at each time point for each person (hereafter, an “allele”) represented a new infection by a parasite strain with that allele (**Methods**). Once probabilities were assigned, alleles were grouped into probability bins (0 to 10%, 10 to 20%, etc., probability of being new), and the proportion of alleles in that bin that were actually new was compared to the bin’s center (average of its endpoints) and weighted by the number of alleles in each bin to compute the probability error.

Without any missing time points, the Bayesian model produced the least biased probabilities for both the rolling presence probability synthetic datasets (1% overestimation versus 4% and 7% underestimation for the simple and clustering models) and the Poisson time to clearance synthetic datasets (3% overestimation versus 11% and 14% underestimation for the simple and clustering models). The clustering model and the simple rule were more likely to mistake repeated observation of a common allele as representing a persistent, rather than a new, infection (**Fig. 2A**). When the proportion of missing time points increased, if the imputation strategy was not used, the predicted probability increased substantially (by 10%-20% over the qPCR range for the models on the rolling presence probability data and 16%-21% for the Poisson time to clearance data) since the missingness-induced gaps appeared as long time periods without previously-observed alleles. However, with the imputation strategy, the predicted probabilities changed much less (1%-5% decrease for any model over the qPCR range), showing that the imputation procedure mitigates bias due to missingness.

**Figure 2.**
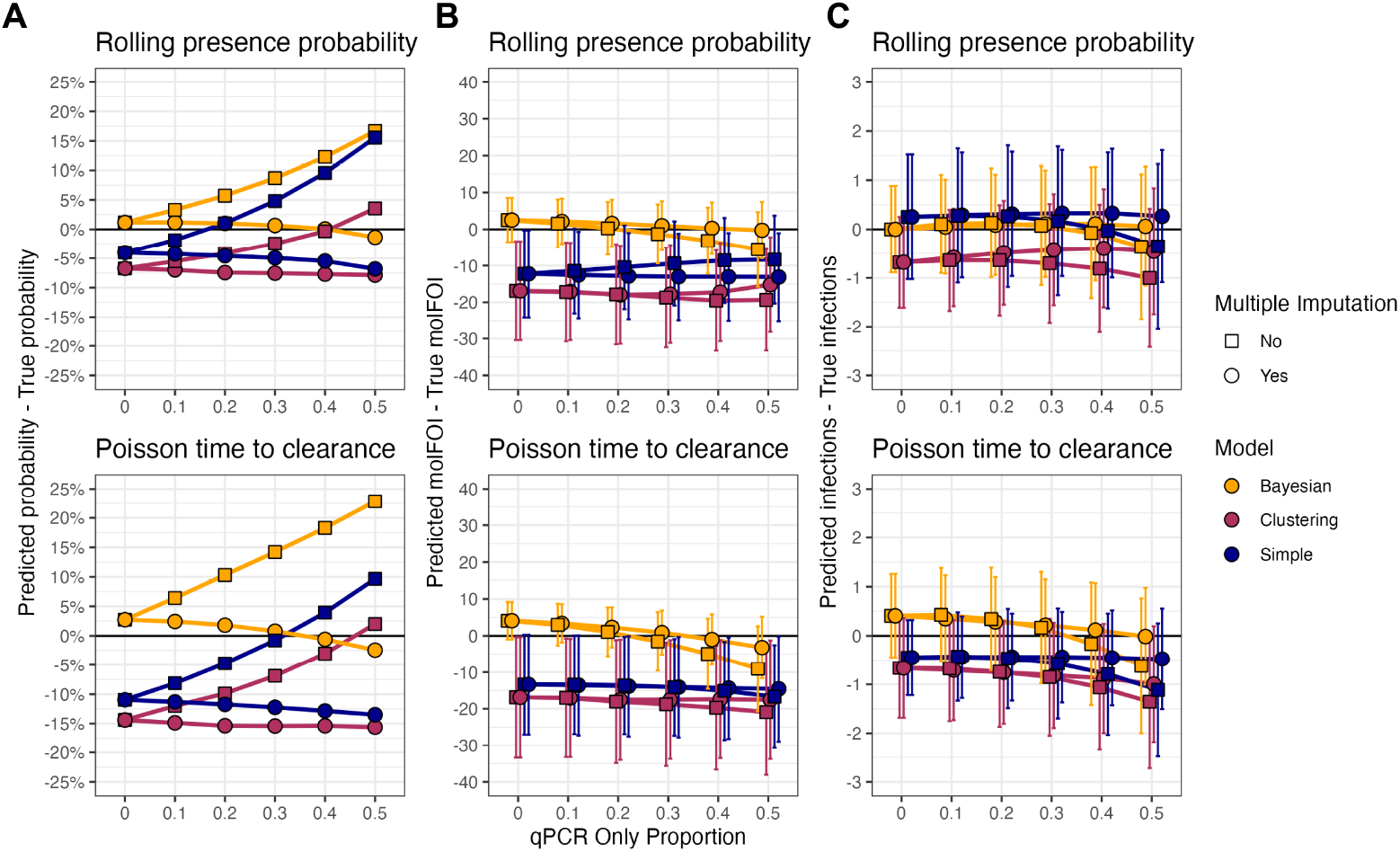
The Bayesian model with the multiple imputation strategy produces approximately unbiased estimates of key infection parameters. Fifty synthetic datasets were generated, each with 200 subjects, an average of 20 evenly spaced time points per subject, one locus with 100 alleles, and varying proportions of sequencing positive samples masked as qPCR-positive only to mimic missingness. New infections were generated according to allele-specific probabilities with non-specific dependencies designed to yield polyclonal infections. Persistent infections were generated according to two data generating models: a “rolling presence probability” model in which the probability an allele was present and persistent was determined at each time point by the allele’s history, and a “Poisson time to clearance” model in which infections were set to have Poisson-length times to clearance. The Bayesian, clustering, and simple models were applied to each dataset, with or without multiple imputation. **A**. Alleles (across all datasets) were binned by their assigned probabilities into intervals of width 10%, and the centers of these bins were compared to the proportion of the alleles in the bin that were actually new, yielding a probability error weighted by the number of alleles in each bin. **B**. The per-subject molFOI was estimated for each subject in each dataset and compared to the true number of new strains (one allele per strain) with which the person was infected across the time points. **C**. The per-subject number of new infection events was estimated for each subject in each dataset and compared to the true number of new infections across the time points. The error mean *±* SD across the subjects and datasets is shown. The probability errors do not have standard deviations because alleles from all subjects and datasets were binned together. Points closer to 0 are better.

The outcomes of interest from longitudinal genotyping studies typically involve metrics such as molFOI and the total number of new infection events per individual over the study period. Therefore, the probabilities assigned by each method were aggregated to estimate these parameters (**Methods**). As before, the Bayesian model produced the least biased estimates of these parameters, and the multiple imputation procedure reduced bias when the dataset contained missing samples (**Fig. 2B-C**). Furthermore, regarding absolute error, the Bayesian model with imputation uniformly estimated the most accurate probabilities, molFOI, and number of new infections across both data types, save that it was slightly less accurate than the simple model at estimating new infections in the Poisson time to clearance data with few missing samples (0.76 vs 0.50 infections from the truth on average with complete data, **Sup. Fig. 1**).

To test the robustness of these results, the number of subjects per datasets, the number of alleles sequenced, and the model properties affecting the rate of new infections were varied (**Supplementary Information**). In general, the Bayesian model was robust to variations in the underlying data and typically produced the most accurate results on all metrics, though the clustering model was the most accurate at estimating the number of new infection events for simulations with either low allele counts (20 and 50) or treatment for infections (**Sup. Figs. 2-3**).

### When using multiple loci, choosing the most variable locus after genotyping maximizes accuracy

To improve new strain detection, it is common for studies to involve genotyping multiple loci instead of a single locus.^9,16,26^ Because alleles at different loci can correspond to the same strain, two de-duplication strategies are implemented to determine the molFOI, which are applied after determining allele-specific probabilities (**Methods, Sup. Fig. 4**). First, the “sum then max” procedure first determines the molFOI for each locus and then chooses the locus with the maximum molFOI for each individual. Second, the “max then sum” procedure first determines the locus with the most new alleles at each time point and then adds these maxima over the time points. As expected, the max then sum method always estimated higher molFOI values, though both methods still underestimated the molFOI when applied after the clustering and simple models (**Sup. Fig. 5**). When applied after the Bayesian model, the max then sum method overestimated the molFOI with any number of loci while the sum then max method estimated correctly with two loci but underestimated with five loci. Still, the sum then max method with the Bayesian model produced the lowest absolute error with one, two, or five loci.

The max then sum method is expected to be least accurate when the loci are sequenced incompletely, spreading the detection of new alleles from a single infection over multiple days. Therefore, the methods were applied to simulations with five loci and varying drop-out rates—the proportion of alleles first sequenced on the second, rather than the first, time point after a new infection. As expected, with higher drop-out rates, the max then sum method increasingly overestimated the molFOI, and its absolute error was only lower than that of the sum then max method when there was no drop-out (**Sup. Fig. 6**). This suggests that, in general, the sum then max method should be used.

### All methods produce highly consistent results on real data, which are frequently unambiguous

To evaluate these methods on real data, three longitudinal malaria genotyping datasets were analyzed: a CIS43LS monoclonal antibody trial cohort,^16^ a 2011 Malian cohort,^9^ and a Ugandan cohort.^5^ The CIS43LS monoclonal antibody trial cohort (348 participants, 5529 time points total) was sampled approximately every two weeks with more frequent sampling in the first weeks of the trial, and samples were analyzed with both qRT-PCR and amplicon sequencing of 6 loci (316 alleles total). The 2011 Malian cohort (648 participants, 5741 time points total) was sampled approximately every two weeks, and samples were analyzed with amplicon sequencing of four loci (548 alleles total) only (i.e., no qPCR). The Ugandan cohort (477 participants, 10884 time points total) was sampled approximately every four weeks for two years, and samples were analyzed with both qPCR and amplicon sequencing of a single locus (45 alleles). Since the CIS43LS trial cohort and Ugandan cohort data included qPCR data (146 of 1014 and 326 of 1187 infection-positive time points were qPCR positive but failed to produce amplicon sequencing data, respectively), multiple imputation was used to create fifty imputed datasets. Then, the Bayesian, clustering, and simple models were run on each dataset (**Methods**). The first two time points for each individual were excluded from analysis because any alleles present could have represented either new or persistent infections. The CIS43LS cohort was characterized by moderate transmission, short infections, and low sequencing drop-out (an estimated 28% of alleles were unobserved when likely present) (**Fig. 3A**). The 2011 Malian cohort was characterized by high transmission, short infections, and high sequencing drop-out (73%) (**Sup. Fig. 7**). The Ugandan cohort was characterized by low transmission, very long infections, and low sequencing drop-out (20%) (**Sup. Fig. 8**).

**Figure 3.**
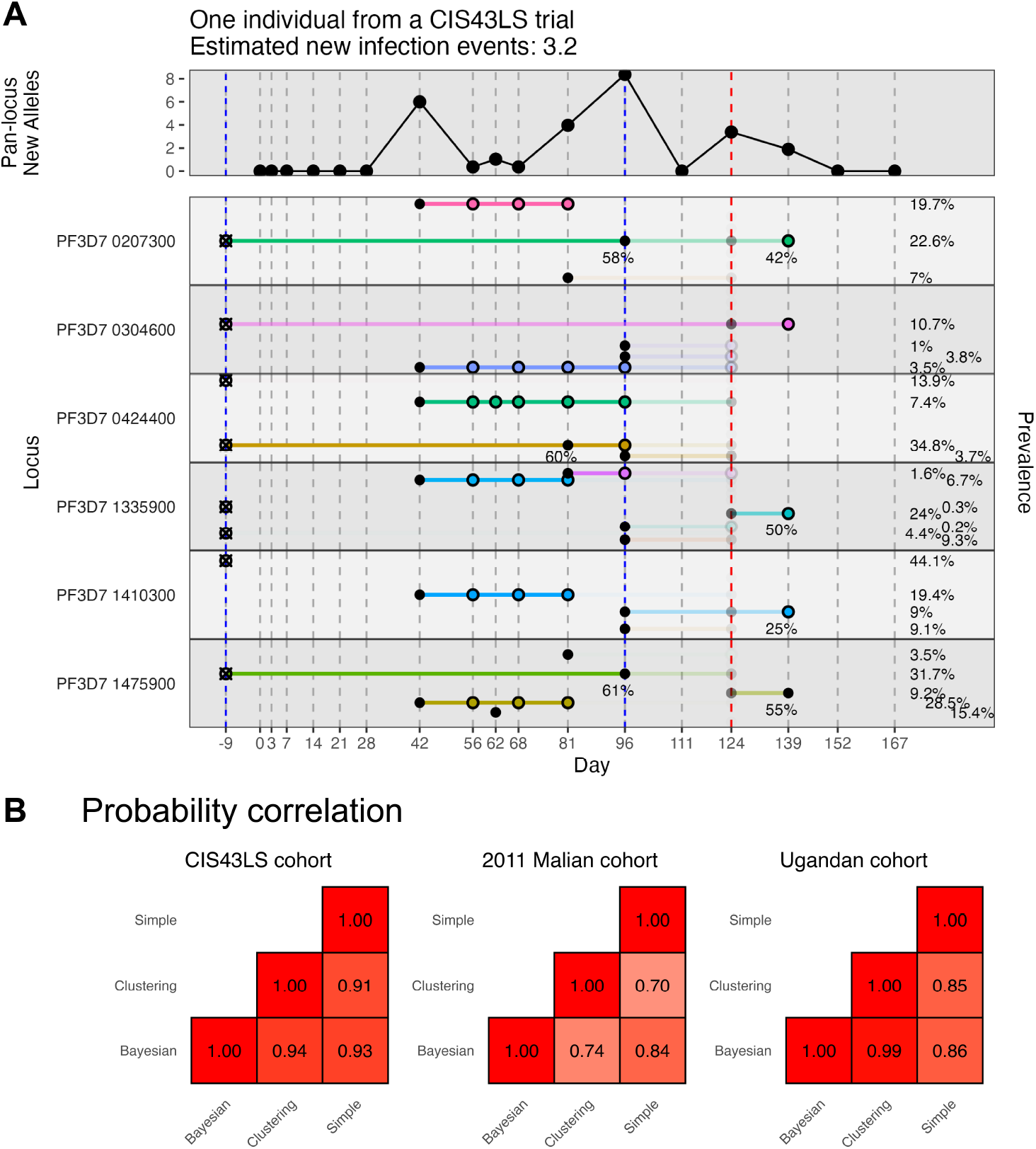
Probabilities assigned to the real datasets are biologically plausible and agree across models. **A**. The default DINEMITES vizualization shows the infection course for the same individual as in **Fig. 1A** with one horizontal pane per locus and one row per allele. Vertical dashed lines mark days when the subject provided blood spots, and dots indicate alleles present in genotyping, connected horizontally if observed repeatedly. Fifty imputed datasets were used, and alleles at missing time points (red dashes, day 124) are opaque proportional to their probability of being present as determined in the imputation process. Treatments for malaria are marked with blue dashes. Probabilities that the alleles were from new infections were assigned using the Bayesian model with parameters for study phase, seasonality, treatment, and previous infections (ever and in the previous 30, 60, and 90 days). Sequenced alleles are marked with solid circles if their average assigned probabilities of being new are over 50%, and they are marked with open circles otherwise. The probability of the allele being new is only displayed under the point if the probability is between 20% and 80%. The pan-locus new alleles (top strip) are calculated by multiplying the probability an allele was present and the probability the allele was new and summing these products over all alleles at each time point. The prevalence on the right is the proportion of sequenced infections in the dataset in which the allele is present. **B**. Correlations between the probabilities assigned by each model on each dataset for present alleles.

The probabilities assigned to each sequenced allele were broadly consistent, with correlations at least 0.70 for all pairs of methods on all datasets (**Fig. 3B**). The probabilities were even more consistent for the CIS43LS cohort and Ugandan cohort, likely due to a combination of more accurate genotyping and the use of the same imputed datasets across methods. Notably, this consistency largely came from the abundance of unambiguous alleles (clearly new or clearly persistent). When subset to alleles assigned probabilities between 5% and 95% by both the Bayesian and clustering methods, the correlations ranged from 0.49 to 0.68 on the CIS43LS cohort data and from 0.14 to 0.55 on the 2011 Malian cohort data compared to 0.91-0.94 and 0.70-0.84 with the full data. When the Ugandan cohort was subset in this way, only 12 of the original 1570 sequenced alleles remained since the alleles from this cohort were almost always categorized as new or persistent with very high confidence.

Next, the probabilities were aggregated to estimate the molFOI and number of infections for each dataset (**Fig. 4A-B**). The results were highly consistent across methods for both the molFOI (correlations 0.92 to 1.00 across the methods and datasets) and the number of new infections (correlations 0.85 to 0.99). The largest differences among the models came from estimating the number of new infections in the Ugandan cohort data since there were frequently instances in which an allele was observed, subsequently not observed for at least three time points, and then observed again (25 instances out of the 126 sequences in which such a pattern could be observed). The simple rule classified these as new infections while the clustering model and the Bayesian model classified these as persistent infections, using the fact that new infections were very rare in this cohort: on average, there were 0.23 new infections per subject with an average of 612 days of observation per subject. Because the 2011 Malian cohort sequencing likely had a high sequencing drop-out rate, a modified Bayesian model was also applied that allowed for drop-out (**Supplementary Information**). As expected, the probability each allele was new, the estimated molFOI, and the estimated number of new infections were all lower with the model allowing drop-out, with the estimated molFOI reduced from 7.9 to 4.9 on average and the estimated number of new infections reduced from 3.2 to 2.1 (**Sup. Fig. 9**).

**Figure 4.**
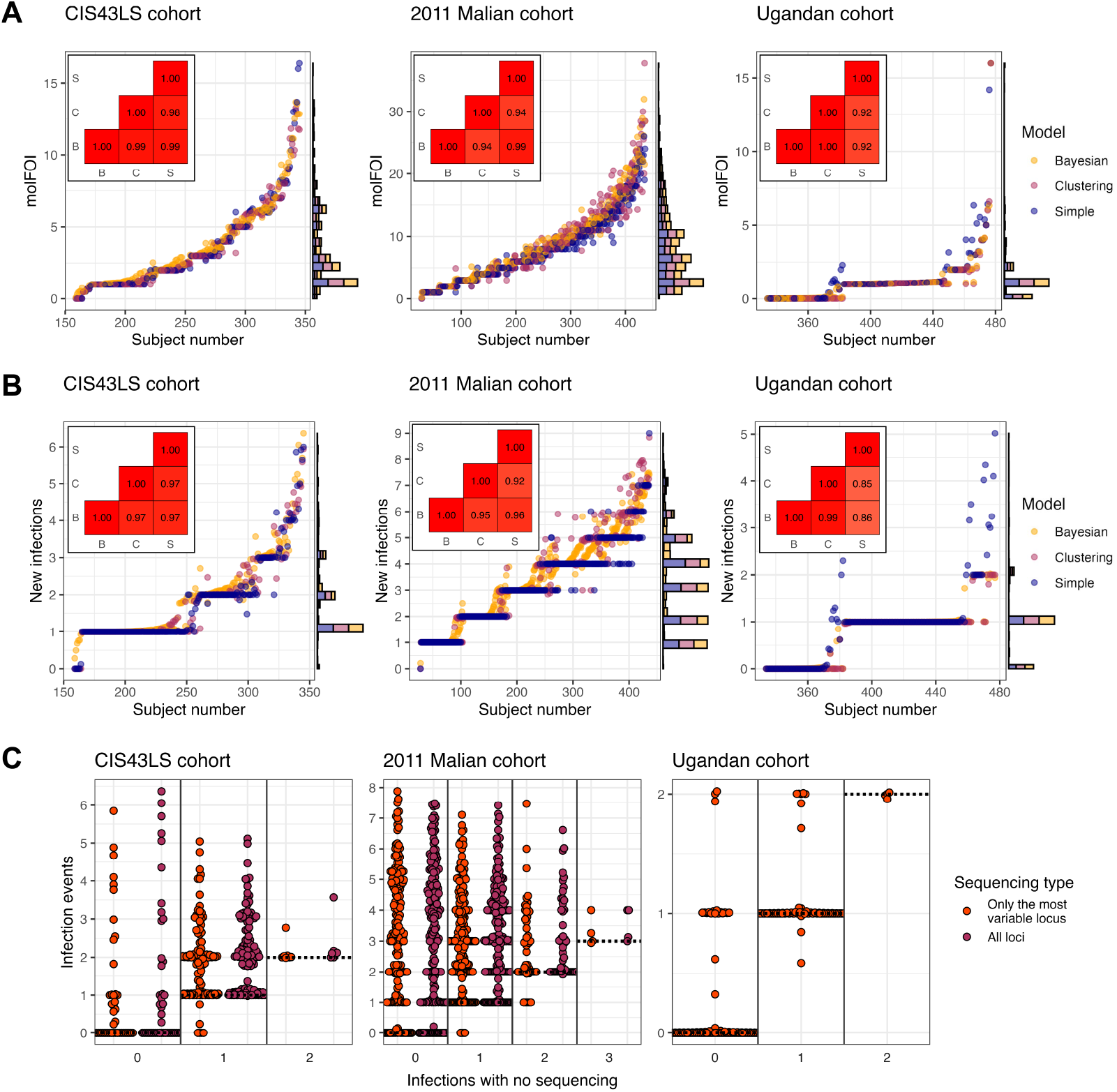
Cumulative infection metrics agreed across models. **A**. Using the sum then max method, the molFOI was estimated for all subjects in each cohort. All subjects not plotted had molFOI values of zero assigned by all models. Insets show the correlations in the estimated molFOI across the Bayesian (B), clustering (C), and simple (S) models. **B**. The number of new infection events was estimated for all subjects in each cohort. All subjects not plotted had no infections estimated by all models. Insets show the correlations in the estimated new infections across the models. **C**. The number of infections predicted by the Bayesian model with all loci sequenced or only the most variable locus sequenced was compared to the count from simply classifying infections as new if no infection had been observed in the previous three time points (i.e., sequencing agnostic).

Finally, to evaluate the combined improvements of sequencing and applying more rigorous statistical methods, the number of infection events was compared between (1) the Bayesian model applied to the full genotyping data or the most variable locus versus (2) a simple rule (the infection is new if not observed in the previous 3 time points) applied to the infection presence/absence data that would have been generated without sequencing. Notably, one strategy might detect no infections while the other detects at least one because the first two time points for each subject were excluded, and subsequent infections might be classified as the same as at those excluded times. Among subjects with at least one infection detected by either the simple rule or genotyping, there were an average of 0.92 infections detected by the simple rule but 1.95 detected by genotyping (112% increase) in the CIS43LS cohort, 0.74 versus 3.45 (359% increase) in the 2011 Malian cohort, and 0.58 versus 0.77 (33% increase) in the Ugandan cohort (**Fig. 4C**). Even when restricting the analysis to only the single locus with the most unique alleles in the CIS43LS and 2011 Malian cohorts, there were 85% and 327% more infections detected on average. Thus, the combination of genotyping and more rigorous statistical methods can substantially improve the detection of new infections.

## Discussion

Longitudinal cohorts sequenced for infectious diseases can answer many biological questions more precisely, but the data they generate can create analytical challenges due to the complicated dependencies between time points, the effect of variable pathogen loads on sequencing success, and uncertainty regarding how best to aggregate allele-specific information into clinical endpoints. Furthermore, previous analyses have relied on plausible but untested assumptions about how long pathogens can remain present but undetected to distinguish new from persistent infections.^5,6,8,20,21^ Distinguishing new from persistent infections is frequently important clinically (e.g., in evaluating interventions that only prevent new infections), but doing so with complete confidence requires expensive and intensive operations such as physically moving people to places they cannot be reinfected.^27^

To address these challenges, we introduced DINEMITES, a set of statistical methods to distinguish new from persistent infections at the allele level and aggregate this information into clinical endpoints including the molFOI and the number of new infection events. On simulated datasets, the Bayesian model produced the most accurate estimates of these parameters across most settings and was well complemented by a multiple imputation procedure to deal with the frequent case of missing time points (**Fig. 2**). When applied to three real datasets, the Bayesian model, the clustering model, and the simple rule implemented in DINEMITES all produced highly consistent estimates of clinical outcomes, bolstering confidence that the results accurately reflect the underlying disease dynamics (**Figs. 3-4**). Finally, when compared to a simple rule applied to only the corresponding infection presence/absence data, the Bayesian model with the complete genotyping data detected 33%, 112%, and 359% more average infections among participants with any infections (**Fig. 4C**).

The high degree of agreement between fundamentally different methods on real data is encouraging because it suggests that the results obtained from such datasets—including previous results—are largely robust to variations in analysis strategies. Indeed, in all datasets, the majority of the sequenced alleles were classified as new or persistent with high confidence by all models. This is expected since every allele observed for the first time must be new, while the same allele observed in short succession is highly likely to be from the same infection, and these two scenarios constitute the majority of genotyping datasets. In fact, the lower accuracy of the non-Bayesian methods in the simulations likely results from the fact that the simulations were constructed to mimic high transmission settings in which each sequenced allele is not as obviously new or persistent.

Practically, these results suggest that future longitudinal genotyping studies should be analyzed by applying all three models (with multiple imputation as necessary for missing time points), comparing the results to check for consistency, and defaulting to the Bayesian model when ambiguous. If multiple loci are sequenced, unless the genotyping is nearly perfect, the recommended method of computing the molFOI is to select the locus with the highest mol-FOI. Finally, multi-locus genotyping is likely still useful for identifying which locus is the most variable, even if sequencing multiple loci did not always substantially increase the number of infections detected compared to sequencing only the most variable locus (**Fig. 4C**). Furthermore, multi-locus genotyping likely improves robustness across *Plasmodium* populations and enables more precise estimates of single time point COI and infection duration.

Despite the typically high accuracy of these recommended methods, there remain some limitations. First, the Bayesian model does not explicitly account for the co-occurrence of alleles, possibly ignoring useful data (though in real data it estimated similarly to the clustering model that uses co-occurrence data). Second, these methods do not incorporate information about read depth or propagate uncertainty in the sequencing results, but rather assume that the genotyping data have been cleaned of errors beforehand and that each allele detected is equally informative. Finally, each of these models still makes important assumptions: the Bayesian model assumes that a prior infection with an allele does not affect the probability of a new infection with that allele; the clustering model assumes that the probably an allele is sequenced depends only on its cluster membership; and the multiple imputation strategy assumes that whether an allele is present at a missing time point depends only on that allele’s presence/absence pattern at nearby time points.

In summary, the methods introduced here represent an important advancement in using more rigorous and reproducible statistical methods to analyze longitudinal genotyping data. While the results here have focused on malaria, these methods could be applied to other infectious disease data in which individuals can clear and reacquire the same pathogen such as tuberculosis, urinary tract infections, rhinovirus, and norovirus.^28,29,30,31,32,33^ These results corroborate the methodology used in previous studies while also providing more rigorous and generalizable methodology for future studies.

## Software and Data Availability

DINEMITES, along with its source code, documentation, tutorials, and example data sets is freely available at https://github.com/WillNickols/dinemites. The code used to run the synthetic and real analyses is available at https://github.com/WillNickols/ dinemites_evaluation. The data used for the Ugandan cohort are available from the corresponding publication,^5^ the raw sequencing data for the 2011 Malian cohort are available from the NCBI Sequence Read Archive under accession PRJNA1129562, and the data used from the CIS43LS and 2011 Malian cohorts are available from the corresponding author upon reasonable request.

## Acknowledgments

The computations in this paper were run in part on the FASRC Cannon cluster supported by the FAS Division of Science Research Computing Group at Harvard University. W.A.N. was supported on a training grant from the National Institute of General Medical Sciences (T32GM135117). This work was also supported by an award from the Bill and Melinda Gates Foundation (INV-052365) and the NIH (1R01AI141544-01). The CIS43LS trial was funded by the Division of Intramural Research and the Vaccine Research Center, National Institute of Allergy and Infectious Diseases, National Institutes of Health. The Mali observational cohort study was funded by the Division of Intramural Research, National Institute of Allergy and Infectious Diseases, National Institutes of Health. We thank Dr. Gail Potter and Dr. Angela Early for informative discussions and feedback on this work.

## Methods

### DINEMITES models

The DINEMITES package implements three models for distinguishing new from persistent infections: a simple model based on time to last occurrence, a Bayesian model based on allele frequencies, and a clustering model based on allele co-occurrence.

### Simple model

The simple model defines an allele as persistent if it has been observed in the last *n*_lag_ visits (default 3) and the last *t*_lag_ days (default infinite). Otherwise, it is defined as new. This captures the intuition that the allele is unlikely to be new if observed in a recent visit, and (optionally) the visit in which it is observed cannot be too long ago.

### Bayesian model

The Bayesian model estimates the probability an allele present represents a new infection by comparing the allele’s background rate of infection to its post-infection probability of being present. The Bayesian model relies on the following key identifiability assumptions:

1. The rate of new infection with an allele is independent of having been infected with a strain with that allele in the past (i.e., no allele-specific immunity)
2. Subjects are independent and exchangeable conditional on their covariates. Notation used for the Bayesian model is summarized in **Table 1**.

**Table 1:**
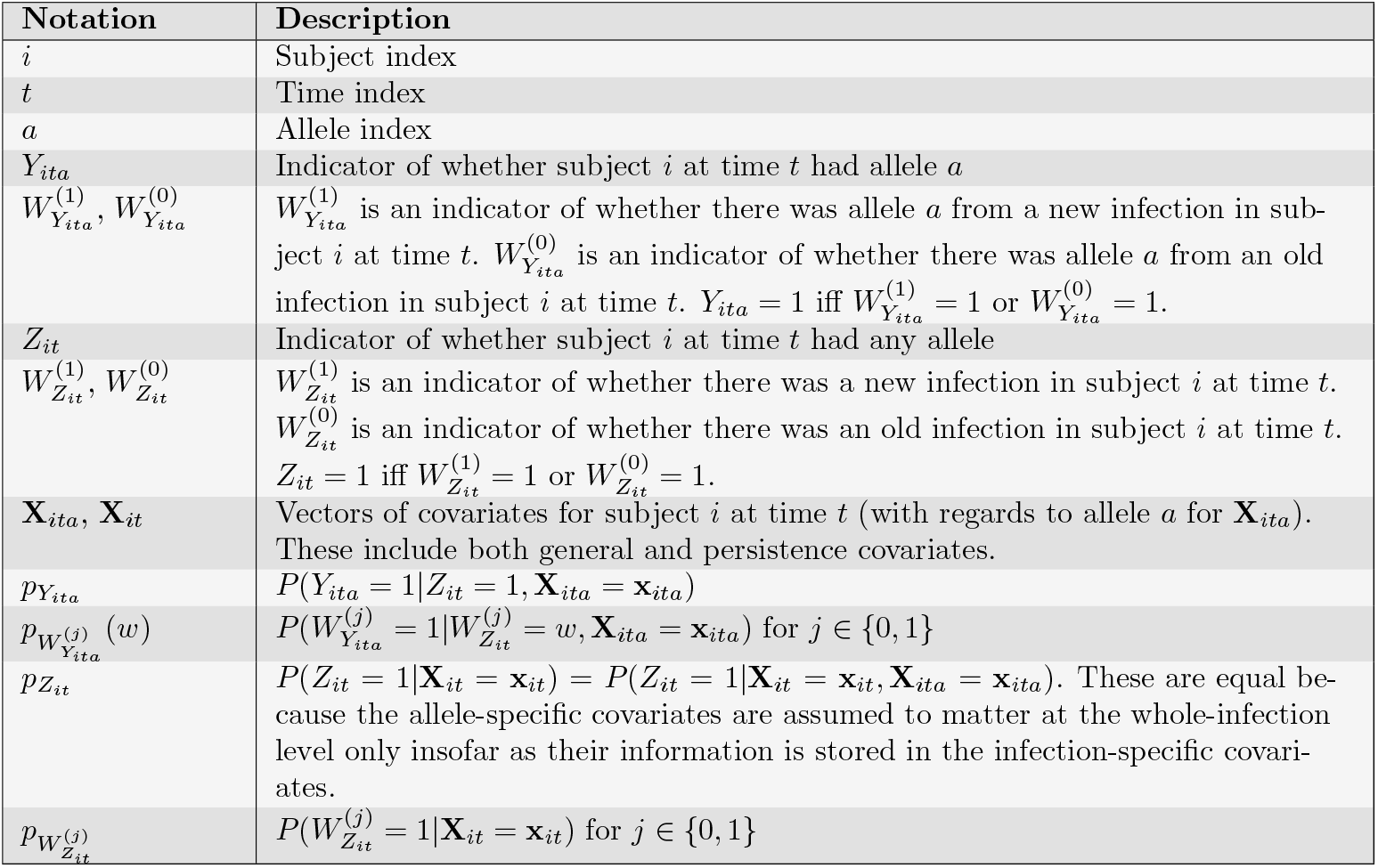
Summary of notation.

Covariates are split into two types: general and persistence. General covariates are those that affect the probability of a new infection (e.g., season or bed net use). Persistence covariates are those that affect the probability of a persistent infection being observed again. These include covariates for if and when an infection has occurred previously along with whether there was treatment between the last observed time and the current time (**Supplementary Information**). Though continuous covariates are allowed, binary covariates allow for deduplication and significant runtime improvements in calculating the likelihood.

The probability there is an infection present for subject *i* at time *t* (any allele) is modeled by a logistic regression with the constraint *β*_*j*_ ≥ 0 if *j* corresponds to a persistence covariate:

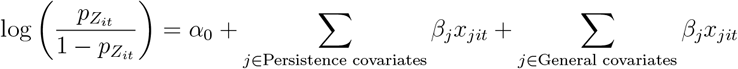

The constraint *β*_*j*_ ≥ 0, using the identifiability assumption, ensures that a previous infection can never decrease the probability of being observed to have an infection. The probability an infection is new is then given by the laws of conditional probability:

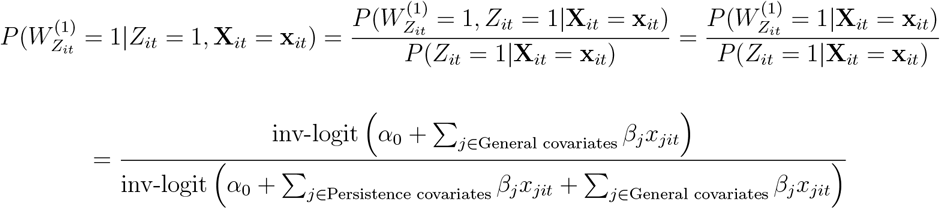

Also, since the new and old infections are assumed to be independent given the covariates, 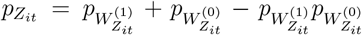. It follows that the probability of an old infection is 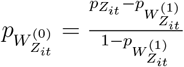.

Next, different logistic models are used for the individual alleles depending on whether the whole infection is new or old:

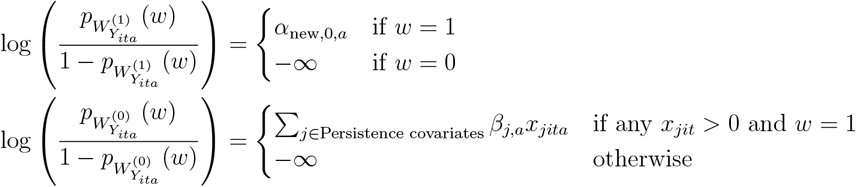

If the infection is new, the probability of an allele being present is just determined by *α*_new,0,*a*_, corresponding to the frequency of the allele in the population. If the infection is old, the probability of an allele being present depends on that allele’s presence/absence history, hence the covariates. However, for the allele to be from an old infection, the infection must be old and the allele must have been present before, hence the log(0) = − ∞. From these parameters, 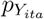 can be calculated (**Supplementary Information**).

The model is parameterized in Stan^34,35^ with *Z*_*it*_ and *Y*_*ita*_ as the observed data. From the fit parameters, the probability each allele is from a new infection can be calculated (**Supplementary Information**). Weakly informative priors for the parameters are specified in the **Supplementary Information**.

By default, the Bayesian model assumes all time points are approximately equally spaced for all subjects or that the probabilities of new infections are unrelated to the length of time between visits. That new infections might be less likely in shorter intervals can be accounted for with general covariates specifying different interval lengths, but modifying the model itself introduces statistical and computational challenges (**Supplementary Information**).

### Clustering model

The clustering model identifies alleles resulting from the same infection through co-occurrence and uses these co-occurrence clusters to estimate whether an allele represents a new infection. The clusters are defined per-subject, corresponding to an assumption of no linkage disequilibrium, which was supported in the data by a lack of cluster similarity across subjects. For each subject, the model does the following:

1. Using the time points at which 2 or more alleles were observed, a graph is constructed with nodes corresponding to alleles and edges with weights given by the number of time points at which the two alleles co-occurred.
2. Using the `getLinkCommunities` function of the `linkcomm` package,^36,37^ edges are hierarchically clustered, and the tree is split at the point that maximizes the partition density to yield sets (clusters) of alleles (possibly overlapping) that represent infection events. Singleton clusters are also added for any alleles that occurred alone.
3. Probabilities are assigned to (1) each cluster occurring on each day and (2) each allele occurring in the clusters to which it has been assigned based on maximizing the likelihood of the observed presence/absence data. Clusters are assumed to be independent of each other.
4. The probability each allele represents a new infection is calculated from these probabilities (**Supplementary Information**).

### Multiple imputation

The multiple imputation procedure takes as input a dataset in which some samples have complete genotyping information and some are only known to be pathogen positive or negative (considered “missing” if positive). The procedure uses the presence/absence patterns of the complete data to generate *n*_imputations_ imputed datasets in which the pathogen positive samples are replaced with complete genotyping information. The multiple imputation procedure does the following *n*_imputations_ times (storing results when redundant):

1. For each subject, for *k* ∈ *{*1, …, *k*_max_*}* (default *k*_max_ = 8), each uninterrupted length *k* sequence of time points (across all alleles) in the complete data is recorded.
2. For each sequence, all possible masks are generated in which all alleles at time points with a positive sequenced infection are masked with missingness indicators.
3. For each allele in the sequence, both the original *k*-ple and masked *k*-ple are recorded, and a table is created recording how often each masked *k*-ple is generated by each complete data *k*-ple.
4. In the dataset, for each time point *t* with missingness, for each allele, the *k*_max_-ple centered at *t* is searched in the table. Then, a complete *k*_max_-ple replacement is sampled from among the complete data *k*-ples that could generate the observed *k*_max_-ple, proportional to their frequencies in the original complete data. Then, the missing value at the center of the *k*_max_-ple is replaced with the value at the center of the *k*_max_-ple replacement, and the window moves to the next time with missingness. If no *k*_max_ples that could generate the missingness pattern are in the original complete data, the search process is repeated with (*k*_max_ − 1)-ples.

Some exclusion criteria are applied to the imputed alleles to ensure they create plausible complete data without distorting downstream metrics (**Supplementary Information**).

The multiple imputation procedure makes the assumptions that whether a missing allele is present or absent depends only on that allele’s presence, absence, or missing pattern at the rest of the time points. To ensure sufficiently many observations are available for the imputation table, no distinction is made between alleles (based on frequency etc.), and time points only matter through their ordering, not their actual values. If visit frequencies are highly irregular, the imputation procedure might not be appropriate.

### Summary statistic calculations

#### molFOI

When only one locus is sequenced, the molFOI is simply estimated by the sum of: for each allele, the probability the allele is new multiplied by the probability the allele is present (1 if sequenced and present, 0 if not present, between 0 and 1 if imputed).

When multiple loci are sequenced, the molFOI aggregated across the loci should not double count pathogen strains. Therefore, two counting procedures are implemented (**Sup. Fig. 4A**). Both start by, for each individual, for each locus at each time point, taking the sum across the alleles of the probability the allele is new multiplied by the probability the allele is present. Then, the two procedures do the following:

1. **Sum then max**: For each locus, sum these sums across all time points. Then, take the maximum of the per-locus sums.
2. **Max then sum**: For each time point, take the maximum of the sums. Then, sum the maxima over the time points.

The first method ensures that drop-out in alleles at different loci at different times does not result in double counting. The second ensures that if different infections have different diversities at the different loci, the different infections are all counted in full.

#### Estimating new infection events

New infection events are identified by finding peaks in the estimated new alleles (spanning multiple times) and then checking the probabilities assigned to the alleles in each set of times representing a peak (**Sup. Fig. 4B**). Peaks (from a local minimum to a local maximum to the next local minimum), rather than singular days with a nonzero number of new alleles, are chosen to represent new infections since typically not all alleles representing an infection are sequenced on the first time point at which the subject has the infection.

The number of new infection events for each subject is estimated as follows:

1. For each time point, compute the number of new alleles (the sum across the alleles and loci of the probability the allele is new multiplied by the probability the allele is present), the maximum probability any allele is new, and the minimum probability any allele is new.
2. Divide the times into peaks demarcated by local minima in the number of new alleles.
3. Within the times belonging to each peak, if the number of new alleles is at least 1, the largest maximum probability is added to the count of new infections. Also, the minimum probabilities for all other time points in the peak are added. If the number of new alleles is less than 1, the peak is assumed to not represent a new infection, and only the maximum of the minimum probabilities at the time points in the peak is added.

When using imputed datasets, the number of new infections is estimated for each subject for each dataset and averaged over the datasets.

### Real data applications

The CIS43LS trial (ClinicalTrials.gov number, NCT04329104) and observational cohort study in Mali (ClinicalTrials.gov number, NCT01322581) were approved by the Ethics Committee of the Faculty of Medicine, Pharmacy and Dentistry at the University of Sciences, Techniques and Technology of Bamako. Written, informed consent was obtained from all study participants including the parents and/or guardians of participating children.

#### Statistical models applied to real datasets

For the CIS43LS trial and Ugandan cohorts, 50-fold multiple imputation was used to generate complete datasets. The models were applied to each imputation, and the probabilities were averaged over the imputed datasets. For the simple model, an allele was considered new if it had not been observed in the previous three time points and persistent otherwise. The clustering model was applied as described. The Bayesian model was applied with infection persistence covariates for whether there had ever been an infection; whether there was an infection in the last 30, 60, and 90 days; and whether there was an acute or longitudinal treatment effect. Equivalent allele-specific infection persistence covariates were also included. For all cohorts, an indicator for the rainy season was used as a general covariate. Also, since the CIS43LS trial involved periods with different visit frequencies, indicators were also used to indicate visits during days 1 to 9 (highest frequency visit phase) or days 10 to 35 (higher frequency visit phase) since new infections were expected to be less likely with less time between visits. Sequencing drop-out rates (the probability of an allele not being observed despite being present) were estimated by computing, for alleles that were observed once and also at either two or three time points later, what proportion of those alleles were also observed at the time points in between. For all models, for each individual, the first two time points were excluded since these could have represented either new or persistent infections.

## Supplementary Information

### Bayesian model

#### Incorporating treatments in the Bayesian model

If an individual was treated, the persistence coefficients for previously observed alleles are returned to 0, but two treatment coefficients are added for those alleles. The first is an acute coefficient set to 1 until the end of the acute treatment phase (default 10 days), and the second is a longitudinal coefficient set to 1 until the next infection with that allele. This captures the intuition that treatment likely clears previous infections (hence setting the other persistence coefficients to 0) but might not (hence the treatment coefficients set to 1).

#### Solving for Bayesian model parameters

The following result will be necessary for solving for 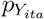. Suppose *A*_1_, *B*_1_, *A*_2_, *B*_2_, *C* are events such that *C* happens if *A*_2_ or *B*_2_ happen, *A*_2_ happens with probability *a*_2_ given *A*_1_ (and 0 otherwise), *B*_2_ happens independently with probability *b*_2_ given *B*_1_ (and 0 otherwise), and *A*_1_ and *B*_1_ happen independently with probability *a*_1_ and *b*_1_ respectively. The probability of interest is *P* (*C*|*A*_1_ ∪ *B*_1_).

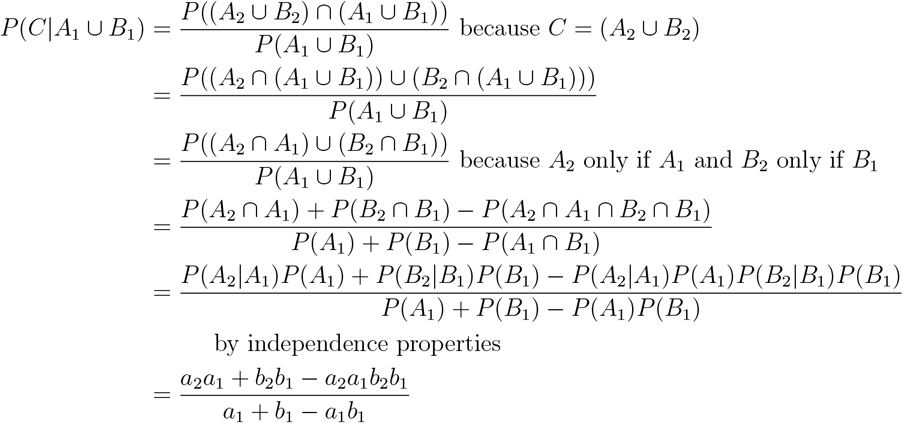

Now, set *Y*_*ita*_ = 1 as *C, A*_2_ and *B*_2_ as 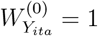 and 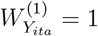 respectively, and *A*_1_ and *B*_1_ as 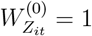 and 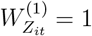 respectively. Using the above,

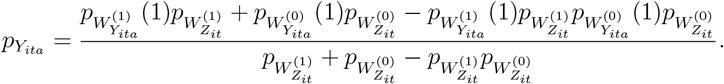

The probability that an allele is new given that it is observed is given by:

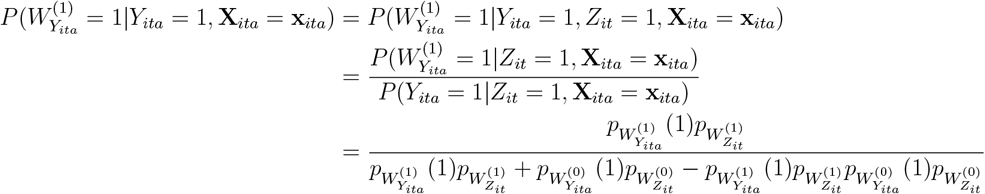

#### Bayesian model with major drop-out

If there is major sequencing drop-out, a model can be used that allows alleles that have never been observed before to be called persistent:

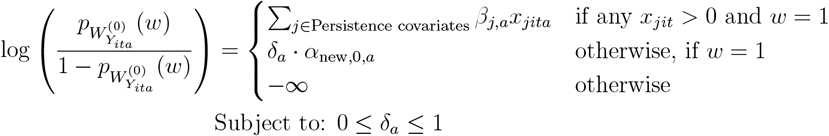

Second, rather than only considering the *Y*_*ita*_ such that *Z*_*it*_ = 1, all *Y*_*ita*_ are modeled, unconditional on *Z*_*it*_ = 1 since the *Z*_*it*_ = 1 are less accurate:

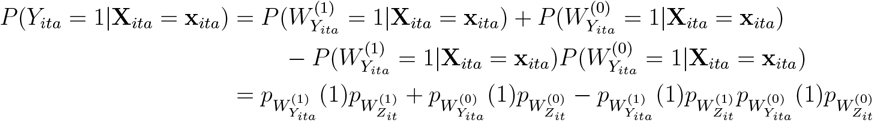

The rest of the model remains the same. In practice, this exposes the second step to many more instances in which *Y*_*ita*_ = 0, leading to lower values for all the parameters and particularly those for new infections.

#### Bayesian model priors

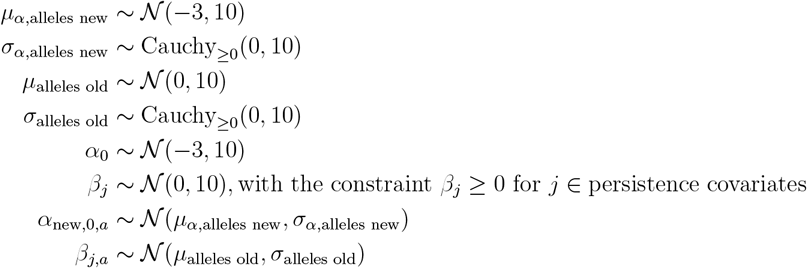

#### Bayesian model specification with major drop-out

When using the drop-out option, the model is augmented by the following:

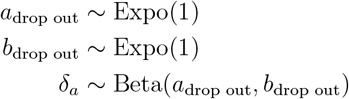

#### Unequal intervals between visits

Because the probability of a new infection in the Bayesian model depends only on the allele and the covariates, shorter times between appointments do not directly result in lower probabilities of new infections. The model could be modified to consider this interval length, but doing so risks inflating the apparent new infection rate if sequencing drop-out at very close time points results in alleles that appear new despite very small intervals. Additionally, this can substantially increase computational complexity because the terms in the likelihood cannot be compressed as effectively when there are many unique interval lengths. If unequal intervals are of concern, general covariates can be included to indicate whether the time point follows e.g. a very short (≤ 2 days), short (3 − 7 days), medium (8 − 21 days), or long (≥ 22 days) interval. This would allow different probabilities of a new infection following each interval, but it could increase variability if there are few intervals of particular lengths.

### Clustering model

#### Probability new calculation

Consider a single subject, and suppose there are *T* time points, *A* alleles, and *N*_*C*_ clusters of alleles 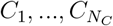 with each cluster being a subset of total alleles (⊆ *{*1, …, *A}*). Let 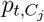 be the probability cluster *C*_*j*_ is present at time point *t*, and let 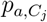 be the probability allele *a* is in cluster *C*_*j*_. Assume these have already been determined from the likelihood maximization step. Then, assuming the clusters are independent of each other,

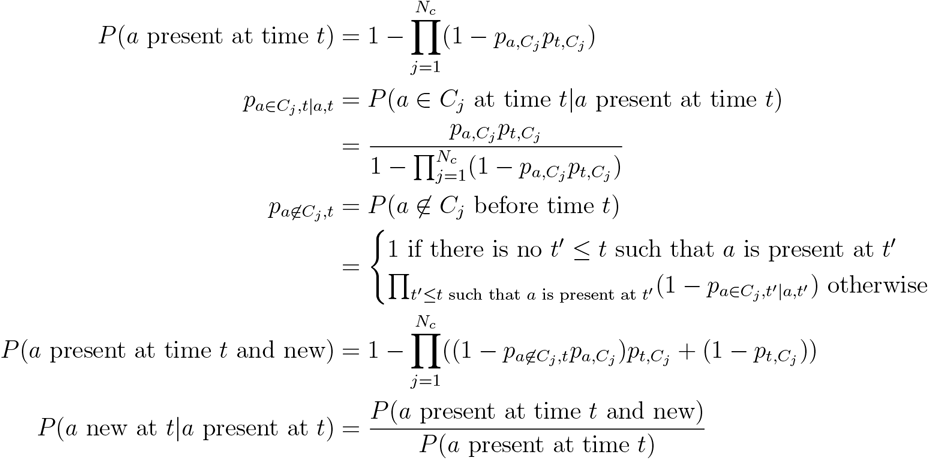

### Multiple imputation criteria

A proposed allele based on the sampling scheme described earlier is only allowed if it satisfies at least one of the following criteria:

1. No arbitrary singletons: the allele has been sequenced or imputed at some other time point for the subject.
2. No spurious infections: the allele is first imputed at the same time point as a second allele is first imputed such that the second allele is sequenced at some time point after being imputed (i.e., the second is not an arbitrary singleton).
3. No neighbors: There are no alleles sequenced within *k*_max_*/*2 time points before or after the current time point.

The no arbitrary singletons and no spurious infections rules together ensure that if there is an infection at a time point with only pathogen positivity, the estimated molFOI remains representatively high without introducing spurious new infection events. The no neighbors rule ensures that if there is effectively no information regarding what allele should be imputed, *some* allele is still imputed to show an infection.

Additionally, imputed datasets are only accepted if all time points with missingness are imputed to have at least one allele present. If this is not the case, the imputation is retried until this is the case or until a retry limit is reached (default 200). If the retry limit is reached, time points are imputed with the closest sequenced allele if the closest is within *k*_max_*/*2 time points and a random allele otherwise.

### Synthetic evaluation data generation

For both synthetic data types (rolling presence probability and Poisson time to clearance), new infections were generated as follows.

1. For each allele (default *n*_alleles_ = 100), a new infection probability is generated from a Beta(1*/*infection multiplier, 50) distribution (default infection multiplier 5).
2. For each subject (default 200 subjects), time points representing appointments (the number of which is sampled uniformly from 7 to 12) are evenly spaced from days 1 to 200.
3. Depending on the model specified, a season indicator is set to 1 for the second half of the time points; a fixed covariate indicator is set to 1 for each subject with probability 1/2; a time-varying covariate is set to 1 at each time with probability 1/2; and/or a prevention covariate indicator is set to 1 for each subject with probability 1/2.
4. Multiplicative modifications to the new infection probabilities are made by (1) sampling from a length *n*_alleles_ multivariate normal centered at 0 with variance 1 and covariance 0.4 (season), variance 0.5 and covariance 0.25 (fixed covariate), or variance 0.5 and covariance 0.15 (time varying covariate); (2) multiplying these modifiers by their indicators; and (3) exponentiating these products and multiplying them by the new infection probabilities.
5. For each subject, at each time point, new infections are sampled as independent Bernoulli random variables with the modified probabilities above. If the number of new alleles is nonzero, a number of additional alleles to be present is sampled from a Poisson with rate parameter equal to the number of new alleles (1) multiplied by the infection multiplier and (2) multiplied by 1/2 if the prevention covariate is 1. Then, that many additional alleles are sampled with probabilities proportional to their modified new infection probabilities.

Finally, new infections are not allowed at neighboring time points since these would be effectively indistinguishable from sequencing drop out in real data. Instead, to simulate sequencing drop-out, with a probability given by the drop-out parameter (default 0.2), each new allele is removed and placed at the next time point as new.

The simulation of persistent alleles differs between the rolling presence probability and Poisson time to clearance models. For the rolling presence probability model, a persistence probability of presence is drawn from a Beta(1, 5) distribution for each allele. Likewise, a lag probability of presence is drawn from a Beta(1, 1) distribution for each allele. Then, on time points after the first infection with an allele, the allele is marked present and persistent according to its persistence probability. If the allele has been observed in the last 30 days, the allele is marked as present and persistent according to its lag probability.

For the models involving treatment, if the subject is infected at the time point, the subject is treated with probability 0.1, and the treatment applies to each allele with a probability of 0.9. If the treatment applies to an allele, the subject is considered to have not had infections from that allele for the purpose of generating persistent infections at subsequent time points. This is intended to simulate treatment that is mostly, but not completely, effective.

For the Poisson time to clearance model, when a set of alleles is new, the number of time points to clearance (for all those alleles) is drawn from a Poisson distribution with rate 2 (80% probability) or a rate equal to the total number of time points (20% probability). These are chosen to simulate short- and long-term infections similar to the lag and persistent infections in the rolling presence probability model.

When there are multiple loci, the alleles are divided evenly among the loci, one locus is chosen as the primary locus at each time point, new alleles are sampled as above at the primary locus, and alleles at each other locus are chosen with one allele per allele at the primary locus (possibly non-unique). In this way, the most diverse locus is selected, and alleles at the other loci are chosen as though there are underlying non-clonal pathogens with particular allele patterns. Then, the Poisson time to clearance model is applied as before.

Finally, for both simulation strategies, time points with sequenced infections are converted to qRT-PCR positive only time points according to the qPCR probability (default 0).

### Synthetic evaluations for robustness

To test the robustness of these results, the number of subjects per datasets, the number of alleles sequenced, and the model properties affecting the rate of new infections were varied. When varying the number of subjects per dataset from 50 to 1000, the accuracy of each model was unchanged (**Sup. Fig. 2**). When varying the number of alleles (but leaving the per-allele infection rates unchanged), the predicted probabilities were mostly unchanged, but the clustering and simple models more severely underestimated the molFOI at high allele counts as increasingly complex infections blurred the distinction between infections. All models underestimated the true number of infections at 200 alleles, but the rate of new infections simulated in that scenario (average 7.5 in 200 days) is likely higher than would be observed in most real datasets. When varying how new infections were generated to account for seasonality, a lack thereof, or seasonality plus a subject-specific protection covariate (e.g., a vaccine or chemoprevention), the accuracy of the models was mostly unchanged. The only infection dynamic to substantially affect accuracy was treatment of subjects for their infections (**Sup. Fig. 2C**). This caused the simple and clustering models to underestimate the probability an allele was from a new infection since these models do not account for treatments which generally eliminate pre-existing infections. It also caused the Bayesian model to overestimate the number of new infections by 0.97 on average (18% above the average 5.4 infections in this high-transmission simulation) despite not affecting the model’s accuracy in determining probabilities or molFOI. Regarding the absolute error, the Bayesian model produced more accurate probabilities and molFOI estimates across all settings, but the clustering model was the most accurate at estimating the number of new infection events for simulations with either low allele counts (20 and 50) or treatment for infections (**Sup. Fig. 3**). Overall, the Bayesian model typically produced the most accurate results and was robust to variations in the underlying data.

### Sequence data generation from blood spots

Each of the three genotyping datasets obtained for this study (from the CIS43LS cohort,^16^ the 2011 Malian cohort,^9^ and the Ugandan cohort^5^) were generated by applying sequential (PCR1 and PCR2) amplification reactions to DNA extracted from dried blood spots collected in Mali and Uganda. In all cases, amplification targeted short *P. falciparum* antigen fragments (188-320 bp, excluding primer binding sites), and sequencing of PCR2-indexed sample pools occurred on an Illumina MiSeq instrument.

For the CIS43LS cohort, PCR amplification involved a multiplex of 6 fragments within the genes CSP (PF37 0304600), TRAP (PF3D7 1335900), SERA8 (PF3D7 0207300), SURFIN (PF3D7 0424400), KELT (PF3D7 1475900), and WD-repeat containing protein (PF3D7 1410300). PCR1 consisted of an initial incubation step at 95°C (3 min); 29 amplification cycles at 98°C (20 s), 57°C (15 s), and 72°C (30 s); and a final extension step at 72°C (1 min). Reaction products subsequently underwent an Exonuclease I digestion and dilution step. PCR2 consisted of an initial incubation step at 95°C (3 min); 10 amplification cycles at 98°C (20 s), 65°C (30s), and 72°C (30 s); and a final extension step at 72°C (1 min). Microhaplotypes were resolved from sequence output by running the malaria amplicon processing pipeline available at https://github.com/broadinstitute/malaria-amplicon-pipeline.git. Alleles were discarded from a sample if represented by *<* 5 read-pairs in the sample or *<* 1% of the total read depth within the sample locus. Alleles which never occurred as major alleles (i.e., never represented the majority of read-pairs within any sample locus) were also excluded from final calls.

For the 2011 Malian cohort,^9^ PCR amplification involved a multiplex of 4 antigen fragments within the genes AMA1 (PF3D7 1133400), SERA2 (PF3D7 0207900), CSP (PF37 0304600), and TRAP (PF3D7 1335900). The latter CSP and TRAP targets are identical to those contained within the six-plex used for the CIS43LS cohort. The PCR protocol used for the 2011 Malian cohort, detailed extensively previously,^26^ is highly similar to that used for the CIS43LS cohort, although it does not involve an intermediate digestion/dilution step. Initial denoising was performed with the same malaria amplicon processing pipeline. Alleles were discarded from a sample if represented by *<* 10 read-pairs in the sample or *<* 1% of the total read depth within the sample locus. Singleton microhaplotypes (i.e., detection in only a single visit) were also excluded from final calls.

For the Ugandan cohort,^5^ PCR amplification involved only a single antigen fragment— the same AMA1 (PF3D7 1133400) target contained within the four-plex used for the 2011 Malian cohort. The PCR protocol used for the Ugandan cohort, previously described,^38^ is slightly distinct in that it uses hemi-nested PCR2 primers to increase sensitivity of the reaction. Also unlike in the CIS43LS and 2011 Malian cohorts, the Uganda workflow processed duplicate samples for each visit, and sequencing output was denoised using SeekDeep methods.^39^

## Supplementary Figures

**Supplementary Figure 1.**
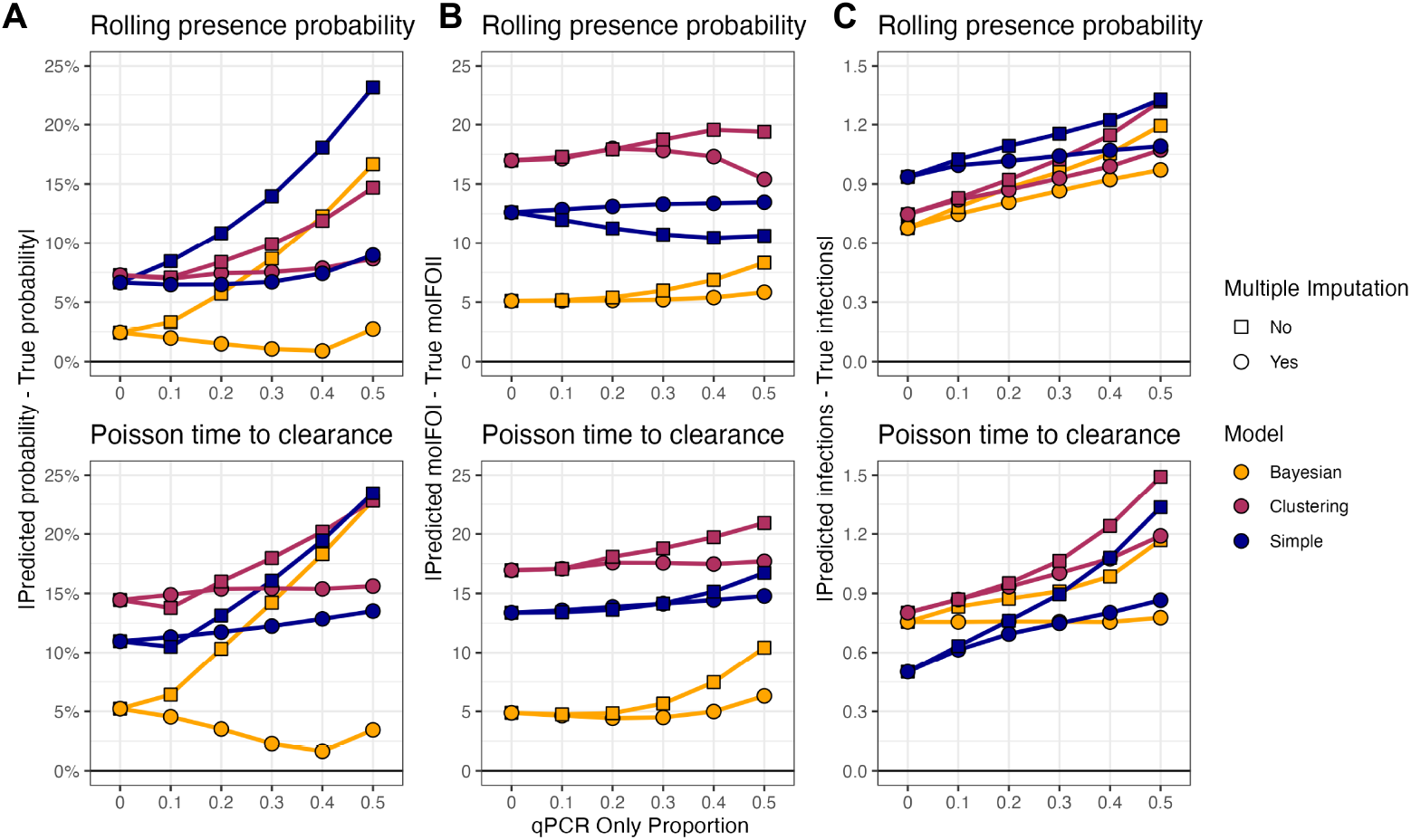
The Bayesian model with multiple imputation typically minimizes the absolute error for the probability an individual sequenced allele was new (A), the per-subject molFOI (B), and the per-subject number of infection events (C). Datasets were generated and error metrics were evaluated as in **Fig. 2**.

**Supplementary Figure 2.**
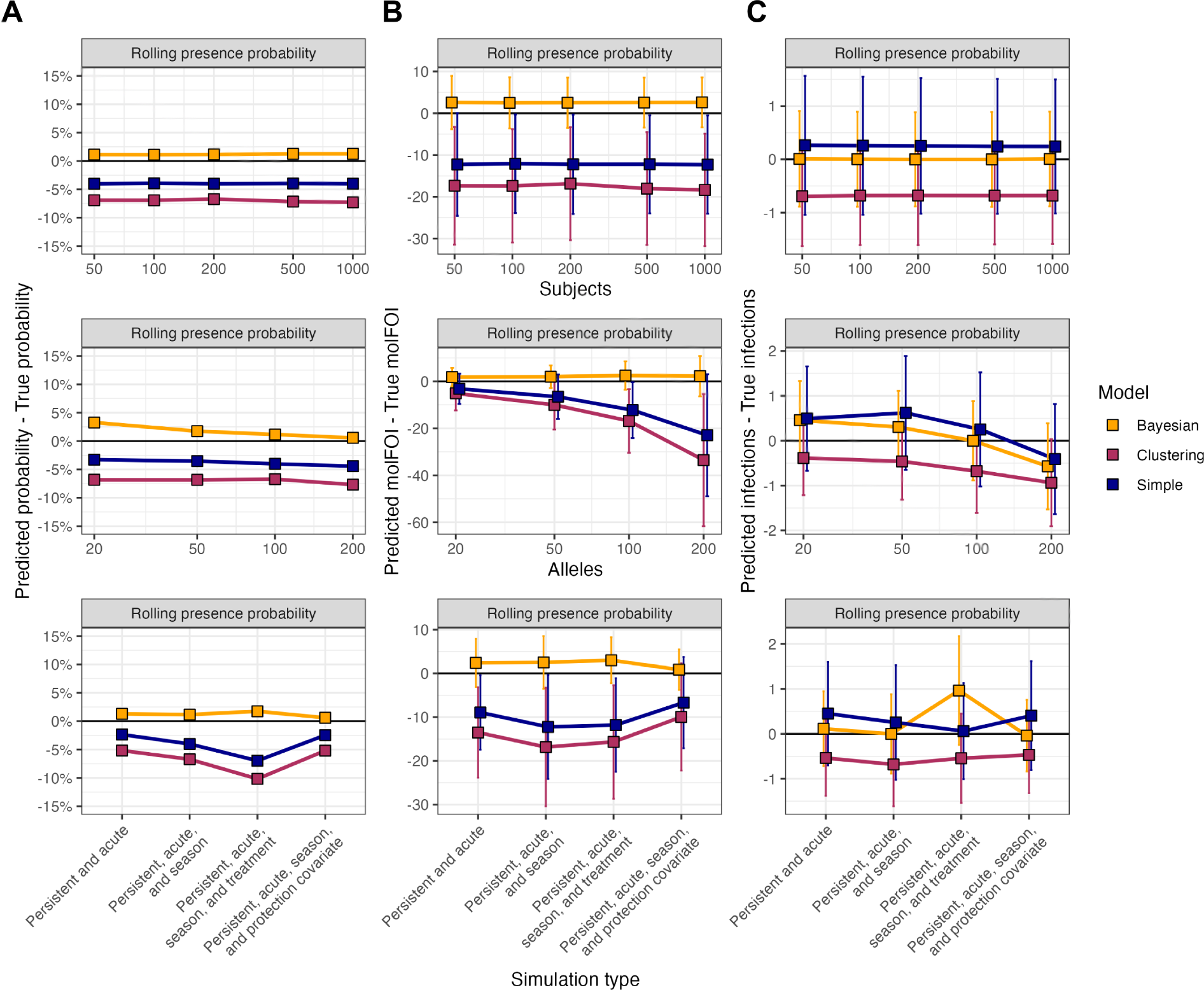
Across different numbers of subjects, different numbers of alleles, and different infection models, the Bayesian model typically produces the least biased estimates of the probability an individual sequenced allele was new (A), the per-subject molFOI (B), and the per-subject number of infection events (C). Datasets were generated and error metrics were evaluated as in **Fig. 2** except with no missing (qPCR only) samples and varying numbers of subjects, varying numbers of alleles, or varying models described in the **Supplementary Information**.

**Supplementary Figure 3.**
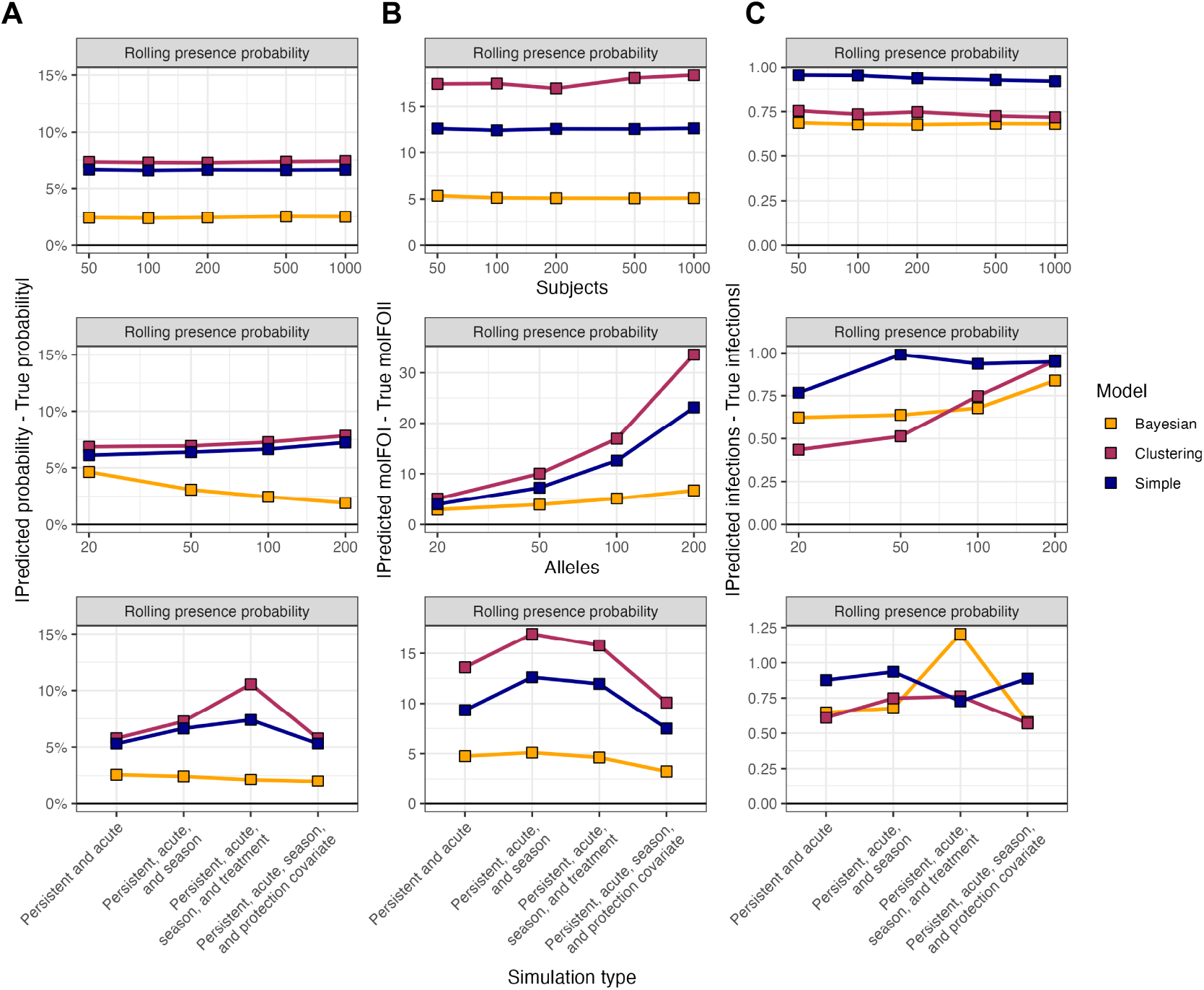
Across different numbers of subjects, different numbers of alleles, and different infection models, the Bayesian model typically produces the lowest absolute error in estimating the probability an individual sequenced allele was new (A), the per-subject molFOI (B), and the per-subject number of infection events (C). Datasets were generated and error metrics were evaluated as in **Fig. 2** except with no missing (qPCR only) samples and varying numbers of subjects, varying numbers of alleles, or varying models described in the **Supplementary Information**.

**Supplementary Figure 4.**
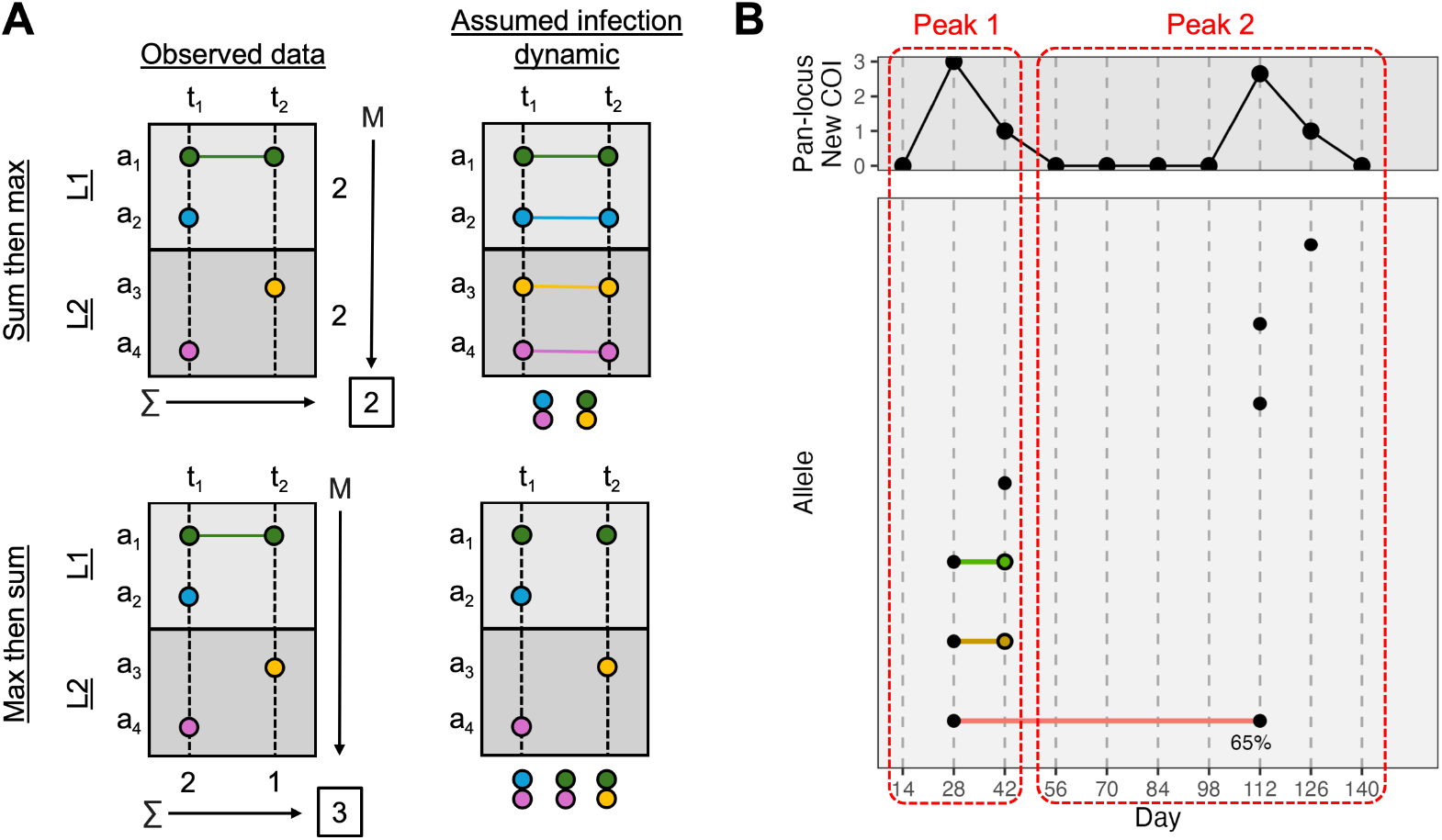
Aggregation methods compute epidemiological endpoints of interest. **A.** For an example time series with four sequenced alleles across two loci, the sum then max and max then sum methods estimate different molFOI values. In the observed data, all alleles are newly observed except *a*_1_ at *t*_2_, which is estimated to be persistent before aggregation. First taking the sum of new alleles for each locus and then taking the maximum of the sums gives a molFOI of 2 while taking the maximum of the number of new alleles per locus per time point and then summing over the time points gives a molFOI of 3. The assumed infection dynamic that would allow each model to be correct is shown to the right. The sum then max model correctly yields a molFOI of 2 when two strains were present at both time points but were sporatically unobserved due to sequencing drop-out. The max then sum model correctly yields a molFOI of 3 when three strains were present at only one time point each and alleles *a*_1_ and *a*_4_ were present in two strains each. For this panel of this figure only, horizontal lines indicate the allele is present due to the same strain persistent at the connected times. **B.** For an example time series, the new infections are estimated. For this panel of this figure only, the solid dots are alleles new with 100% probability unless otherwise labeled, and the open dots are alleles new with 0% probability (i.e., persistent). The time series is divided into two peaks of pan-locus new alleles, each of which contains a time point with an allele with a 100% probability of being new (days 28 and 112), yielding an initial count of 2 new infections. Additionally, the minimum probability an allele is new from each other time point is added. This adds zero at day 42 since the green and yellow alleles have probability zero of being new but adds 1 at day 126 since the top allele is the only one present and has a 100% probability of being new. Thus, the estimated new infection count is 3.

**Supplementary Figure 5.**
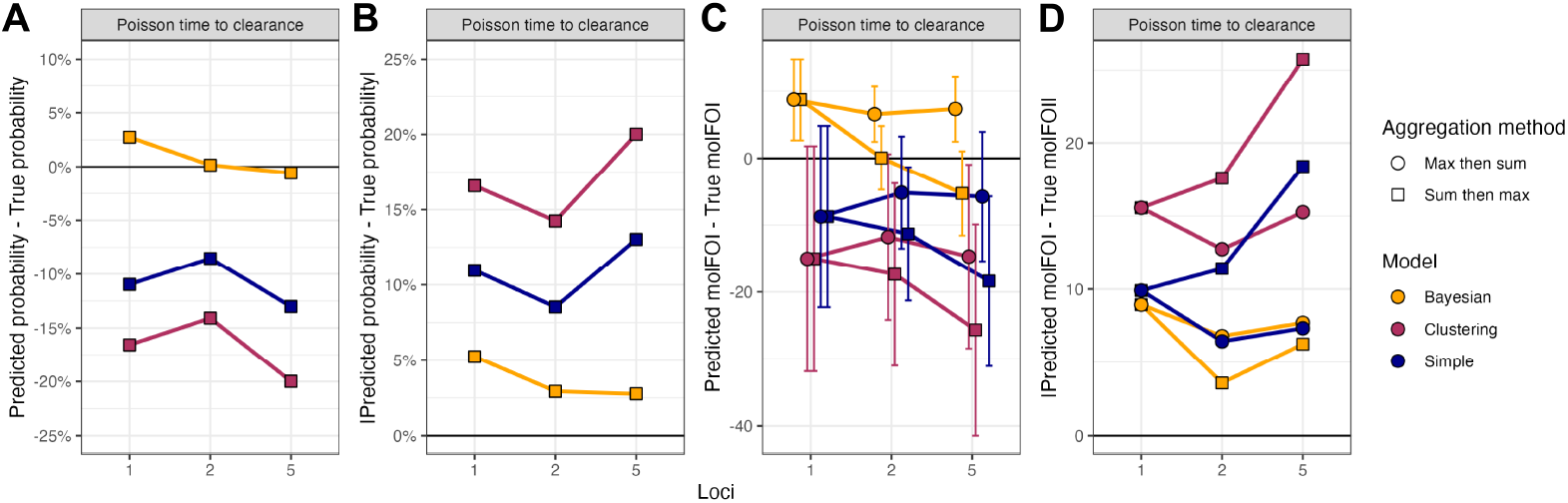
Across different numbers of loci sequenced, the Bayesian model typically produces the least biased and lowest absolute error estimates of the probability an individual sequenced allele was new and the per-subject total molFOI. Datasets were generated and error metrics were evaluated as in **Fig. 2** except with no missing (qPCR only) samples and varying numbers of loci with alleles generated as described in the **Supplementary Information.** For computing the molFOI, both the “max then sum” and “sum then max” strategies were applied. New infections were not evaluated because alleles across loci are treated the same as alleles within a locus for determining new infections.

**Supplementary Figure 6.**
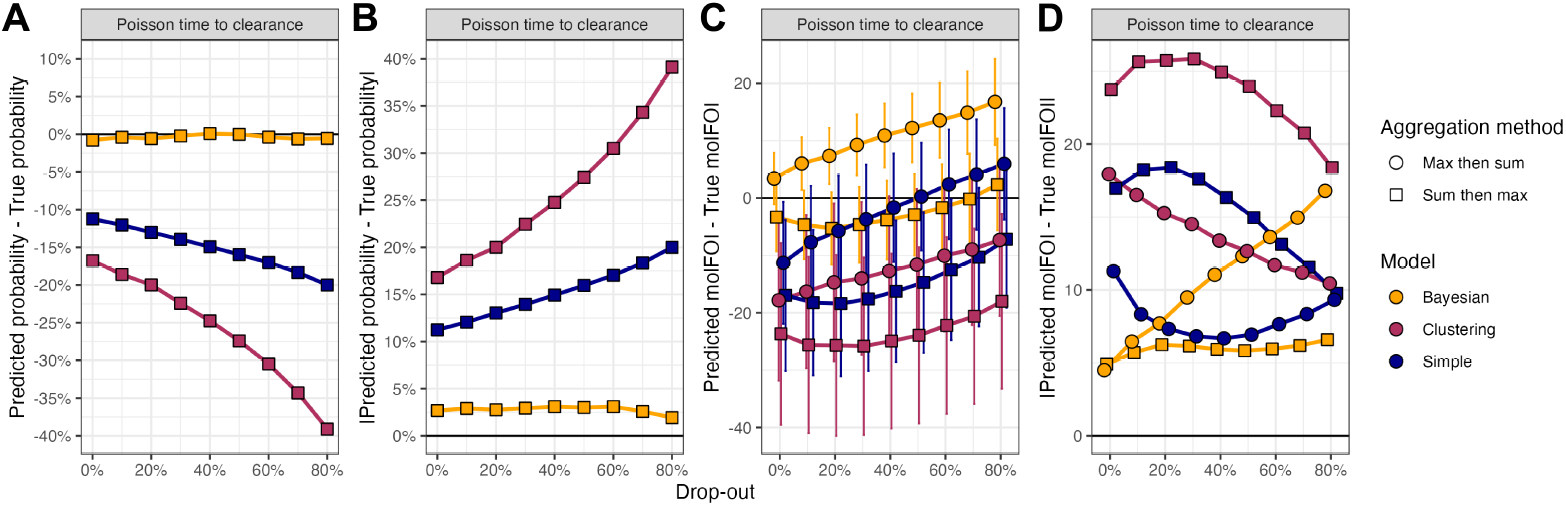
Across different rates of sequencing drop-out at the first time point of an infection, the Bayesian model typically produces the least biased and lowest absolute error estimates of the probability an individual sequenced allele was new and the per-subject molFOI. Datasets were generated and error metrics were evaluated as in **Fig. 2** except with no missing (qPCR only) samples and five loci with alleles generated as described in the **Supplementary Information.** For computing the molFOI, both the “max then sum” and “sum then max” strategies were applied. New infections were not evaluated because alleles across loci are treated the same as alleles within a locus for determining new infections.

**Supplementary Figure 7.**
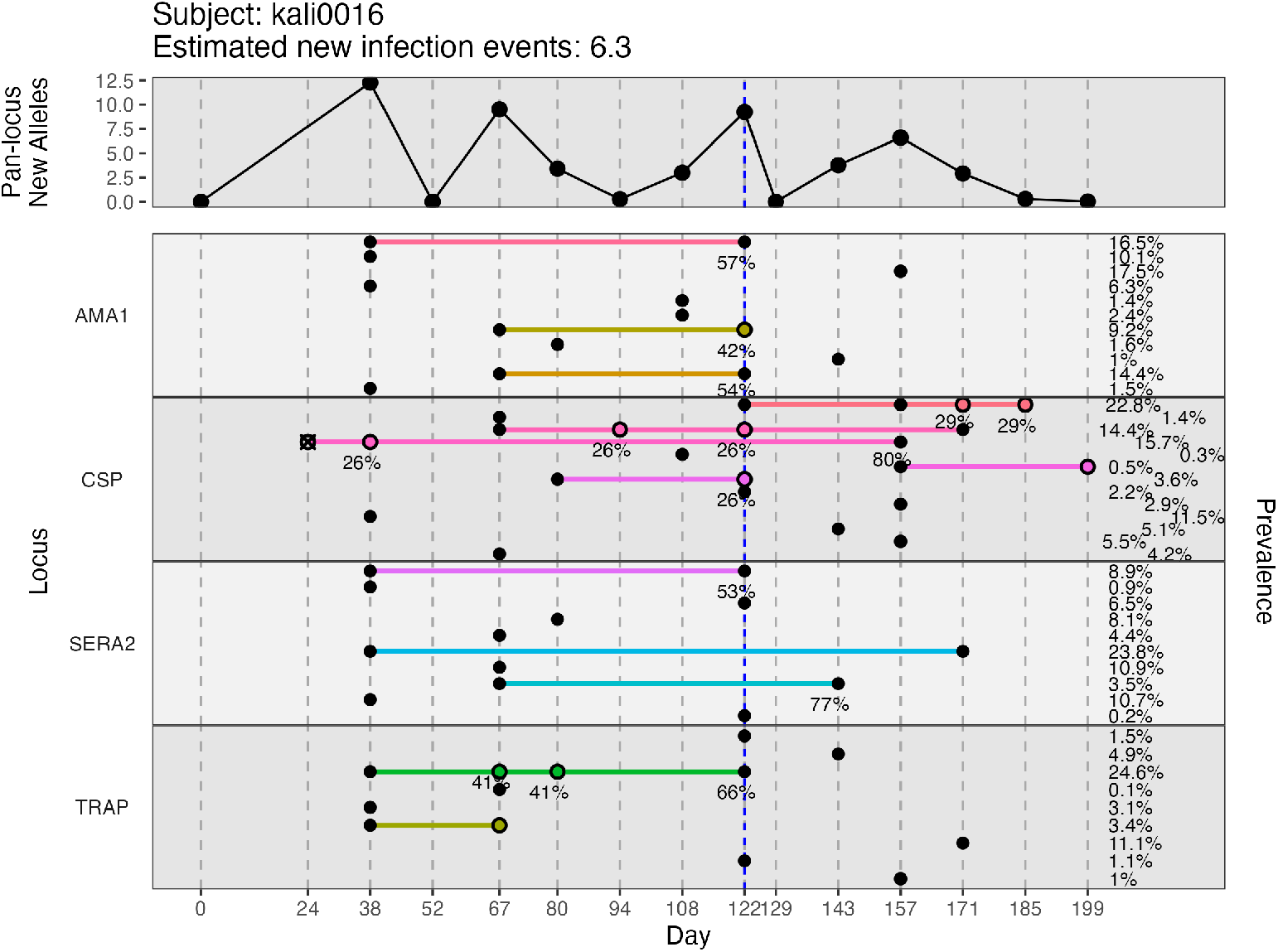
A representative example of longitudinal data from an individual in the 2011 Malian cohort assigned probabilities with the Bayesian model. The default DINEMITES vizualization shows the infection course for one individual with one horizontal pane per locus and one row per allele. Vertical dashed lines mark days when the subject provided blood spots, and dots indicate alleles present in genotyping, connected horizontally if observed repeatedly. Treatments for malaria are marked with blue dashes. Probabilities that the alleles were from new infections were assigned using the Bayesian model with parameters for seasonality, treatment, and previous infections ever and in the previous 30, 60, and 90 days. Sequenced alleles are marked with solid circles if their average assigned probabilities of being new are over 50%, and they are marked with open circles otherwise. The probability of the allele being new is only displayed under the point if the probability is between 20% and 80%. The pan-locus new alleles is the sum of the probabilities over all alleles at each time point. The prevalence on the right is the proportion of sequenced infections in which the allele is present.

**Supplementary Figure 8.**
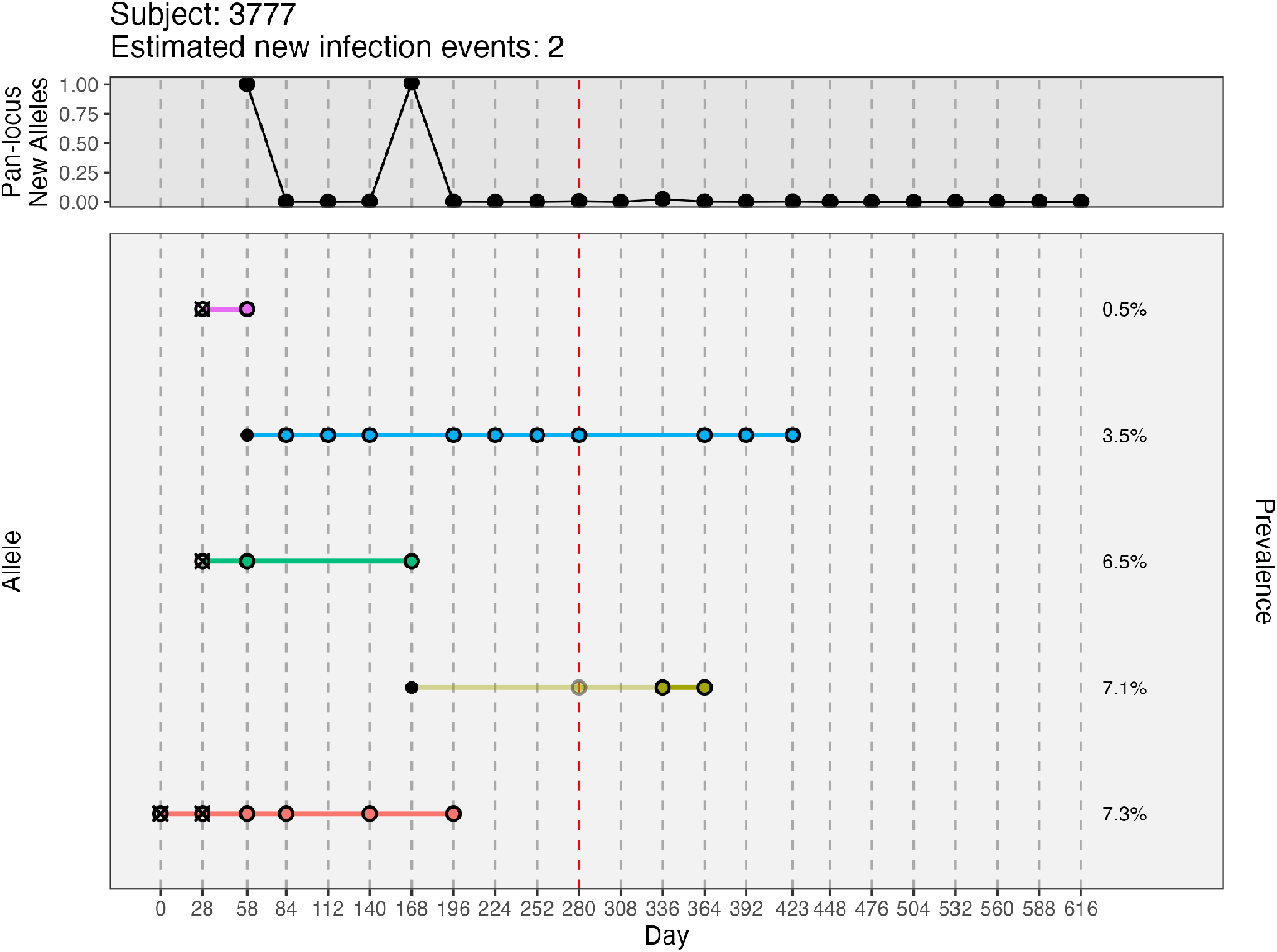
A representative example of longitudinal data from an individual in the Ugandan cohort assigned probabilities with the Bayesian model. The default DINEMITES vizualization shows the infection course for one individual with one horizontal pane per locus and one row per allele. Vertical dashed lines mark days when the subject provided blood spots, and dots indicate alleles present in genotyping, connected horizontally if observed repeatedly. Fifty imputed datasets were used, and alleles at missing time points (red dashes) are opaque proportional to their probability of being present as determined in the imputation process. Probabilities that the alleles were from new infections were assigned using the Bayesian model with parameters for seasonality, treatment, and previous infections ever and in the previous 30, 60, and 90 days. Sequenced alleles are marked with solid circles if their average assigned probabilities of being new are over 50%, and they are marked with open circles otherwise. The probability of the allele being new is only displayed under the point if the probability is between 20% and 80%. The pan-locus new alleles is the sum of the probabilities over all alleles at each time point. The prevalence on the right is the proportion of sequenced infections in which the allele is present.

**Supplementary Figure 9.**
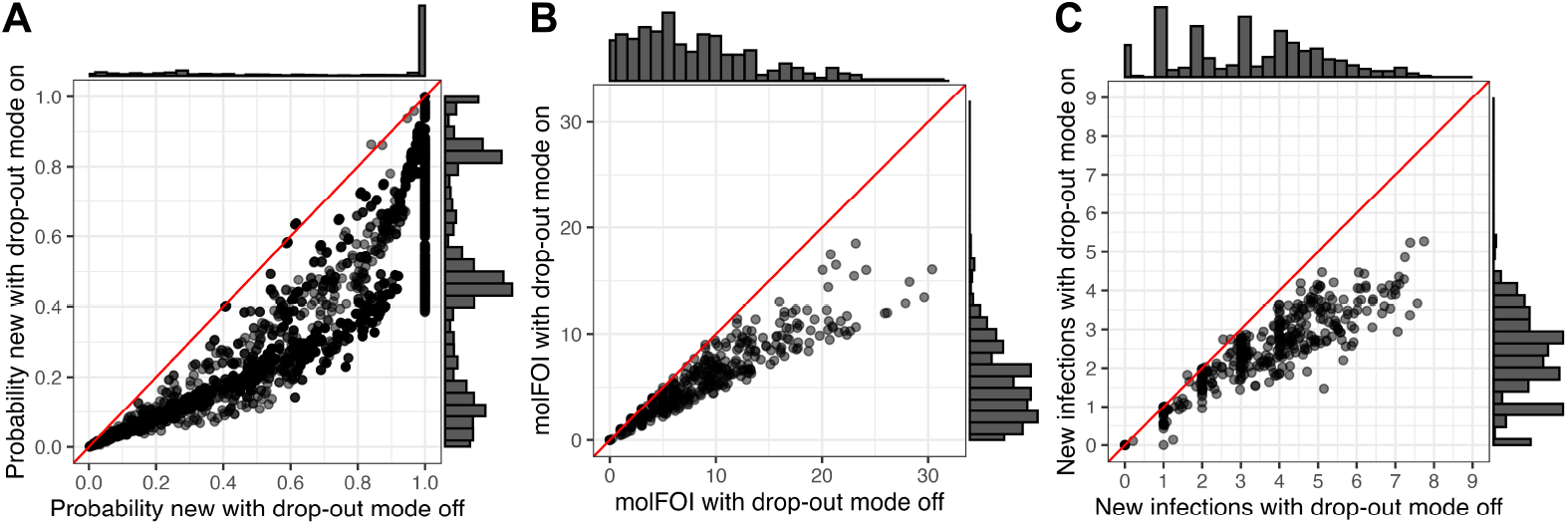
A Bayesian model allowing for newly observed alleles to not be counted as new due to sequencing drop out produced lower estimates of the probability an individual sequenced allele was new (A), the per-subject molFOI (B), and the per-subject number of infection events (C). For the 2011 Malian cohort, the same model parameters as described in the **Methods** were used but with the drop-out model described in the **Supplementary Information**.

